# Fundamental steps in Minichromosome Maintenance complex assembly are conserved from archaea to eukaryotes

**DOI:** 10.1101/2023.08.11.552962

**Authors:** Oliver W Noble, Clement Degut, Michael R Hodgkinson, James P J Chong, Michael J Plevin

## Abstract

Minichromosome maintenance (MCM) proteins are the replicative helicase in both archaea and eukaryotes. Recent cryo-EM studies have defined key steps in the assembly pathway of the eukaryotic MCM2-7 complex and determined roles for each subunit. While archaeal MCMs are simpler in composition, we know little about how these homohexameric MCMs assemble. Investigation of archaeal MCMs has largely focussed on proteins from thermophilic organisms, which typically form robust oligomers under ambient experimental conditions, preventing dissection of their assembly pathway. Here, we describe an uncharacterised MCM from the mesophilic archaeon *Mancarchaeum acidophilum* (*Mac*MCM). A 3D structure of *Mac*MCM reveals subunit-subunit interfaces that are more similar to yeast MCM2-7 than previously studied MCMs from thermophilic archaea. We show that assembly of a *Mac*MCM homohexamer on DNA proceeds via comparable steps to MCM2-7. These results reveal an ancient mechanism underlies assembly of the MCM complex, which is conserved from archaea to eukaryotes.

## Introduction

DNA replication is a fundamental process carried out by all living organisms. At the core of the replication machinery is a processive hexameric helicase that catalyses strand separation of parental double-stranded (ds) DNA ahead of a DNA polymerase^1^. Across the domains of life, two core replicative helicase families have evolved: archaea and eukaryotes utilise minichromosome maintenance (MCM) helicases^2^, while bacteria use DnaB^3^.

The MCM complex was initially discovered in yeast^2^. Six closely related genes, MCM2-7, were identified which were later demonstrated to assemble into a ring-shaped heterohexameric complex with defined subunit order^4^. Each subunit in the MCM2-7 heterohexamer has evolved distinct roles that fine tune regulation of both complex assembly and DNA unwinding. MCM2 and MCM5 are neighbours in the ring. The transient interaction between these subunits is functionally critical and creates a defined opening in MCM2-7 ring known as the MCM2/5 gate^5^. *In vitro*, closing of this gate is a slow step that is dependent on ATP hydrolysis and which precedes stable association of the full MCM2-7 complex with DNA^6^.

In recent years, cryo-EM has provided significant insight into how the eukaryotic MCM2-7 heterohexamer loads onto a DNA substrate and the role of protein cofactors in this process^7^. The stabilities of the different states in the assembly pathway are dependent on cofactors and experimental conditions including temperature, salt and sample concentration^8^. A key point in the assembly of a functional MCM2-7 involves an open ring state, which is stabilised by interaction with ATP and the protein cofactor Cdt1^9^. In this conformation, the C-terminal winged helix domain (WHD) of MCM5 is found in the central channel of the heterohexamer, where it prevents the interaction of MCM5 with MCM2 and therefore closure of the ring^10^. Cdc6 and the Origin Recognition Complex (ORC) recruit open-form Cdt1-MCM2-7 by interacting with the WHDs of MCM2-7^11^. DNA is then threaded into the open MCM2-7 ring, displacing the WHD of MCM5 from the central channel and allowing closure of the MCM2/5 gate^11^. Hydrolysis of bound ATP by MCM2-7 results in release of Cdt1 and stabilisation of a closed DNA-bound MCM2-7 hexamer^8,11^.

Unlike eukaryotes, most archaeal genomes carry only a single MCM gene ^5^. The resulting homomeric composition of archaeal MCMs and the capacity to produce them recombinantly using *Escherichia coli* has meant that much of our early appreciation of the core structural biology of MCMs was obtained from studies of archaeal proteins. However, while proving beneficial in many regards, the homomeric composition of archaeal MCMs has also complicated fundamental questions about MCM structural biology and evolution.

The centrality of the MCM2/5 gate in the loading of a MCM2-7 complex onto DNA makes it difficult to rationalise how related the assembly mechanism of a homohexameric MCM on DNA is given that it has six equivalent subunits. Here, we propose the early focus on characterising MCMs from thermophilic archaea has limited our understanding of homomeric MCMs. All previous *in vitro* studies of homohexameric systems have been conducted using archaeal MCMs that predominantly form oligomers in the absence of ligands such as ATP or DNA. Most techniques that are used to assess oligomeric state are limited to conditions approaching room-temperature, well below the temperatures that enzymes from thermophilic organisms would experience *in vivo*. Taken together, it has been challenging to investigate how homohexameric MCM complexes assemble and hence to understand whether mechanistic similarities between eukaryotes and archaea exist.

We therefore sought to expand our understanding of archaeal MCMs by investigating a selection of previously unstudied examples from species that inhabit a broader selection of environmental niches. This allowed us to identify an MCM from *Mancarchaeum acidiphilum* (*Mac*MCM) that has robust helicase activity at room temperature. We elucidated the 3D structure of *Mac*MCM and determined that its properties in solution are more similar to heterohexameric eukaryotic MCMs than previously characterised homohexameric MCMs from thermophilic archaea. The discovery of *Mac*MCM permitted us to examine the assembly of a homohexameric MCM complex for the first time, and to identify steps in this process that are fundamental and conserved across MCMs.

## Results

### An archaeal MCMs with robust activity at ambient temperature

Historically, over 95 % of studies of archaeal MCMs have focussed on enzymes from organisms that occupy high temperature environments (>65 °C; Figure 1a). To evaluate whether our understanding of the biochemistry of archaeal MCMs has been biased, we sought to assess the activity of MCMs from a broader range of archaea (Figure 1b). We chose three well studied MCMs from thermophilic organisms for which structures have been reported (*Mth*MCM, *Sso*MCM and *Pfu*MCM); five MCMs from thermophilic archaea which live in high temperature environments or which represent more distant phylogenetic lineages (*Ape*MCM, *Afu*MCM, *Kcr*MCM, *Mka*MCM, *Neq*MCM) (Figure 1b-c); and six MCMs from the genomes of mesophilic archaea (20-45 °C), which are adapted for life in various distinguishable habitats, including saline (*Nma*MCM), hypersaline (*Hvo*MCM*, Nac*MCM, *Mha*MCM), anaerobic (*Mba*MCM) and acidic (*Mac*MCM). Of these 14, three are encoded by the genomes of parasitic/symbiont archaea (*Nac*MCM, *Neq*MCM, *Mac*MCM), all of which have extremely small genomes (<1 Mb). Bioinformatic analyses were performed on each sequence to confirm the presence of conserved subdomains and motifs within each MCM (Supplementary Figure 1-2)^12–14^. To conclude our selection, we included a previously engineered chimeric MCM, generated by fusing the N-terminal domain (NTD) of *Sso*MCM with the ATPase domain of *Pfu*MCM (*Sso_N_Pfu_C_*MCM)^15^, as well as the reverse chimera, *Pfu_N_Sso_C_*MCM. The regulatory winged helix domain (WHD) was not included in either chimeric enzyme.

**Figure 1.**
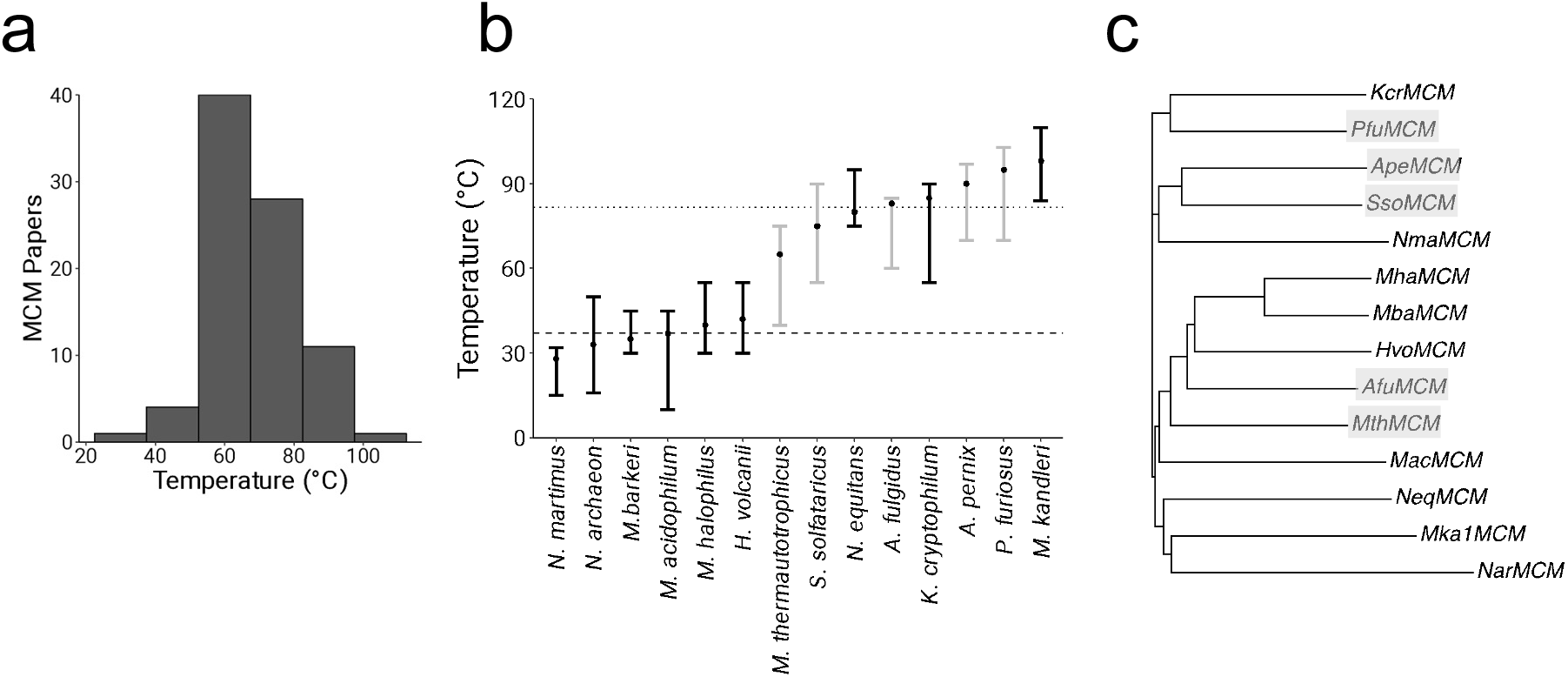
Selection of a diverse library of 14 archaeal MCMs. **a** Comparison of the number of publications released on archaeal MCMs against the natural temperature experienced by the enzyme in vivo. **b** Comparison of the preferred environmental temperatures of the organisms targeted. Grey, enzyme has been previously characterized; black, uncharacterized. Data points represent optimal growth temperature of organism; bars represent growth range. See Supplementary Table 1 for supporting references. **c** Phylogram based on the sequence alignment of the 14 naturally occurring MCM sequences studied here. Sequences were retrieved from the KEGG database^13^ and a phylogenetic tree constructed using Clustal Omega^14^.

Recombinant His_10_-MCMs were overexpressed in *E. coli* and protein expression and solubility assessed using gel electrophoresis (Supplementary Figure 3). Bands at molecular weights (MW) consistent with at least 11 of the MCM targets were observed in the soluble fraction following centrifugation of cell lysates. In general, MCMs from thermophilic organisms expressed at higher levels and were more soluble than MCMs from mesophilic organisms. Each MCM was purified using a single immobilised metal affinity chromatography step. Different ranges of sample purity and nucleic acid contamination were observed for each MCM construct (Supplementary Figure 4; Supplementary Table 2).

A fluorescence-based helicase assay^16^ was used to assess the dsDNA unwinding activity of each sample (Figure 2a). For comparison, unwinding values were standardised to protein concentration (unwinding % per 1000 nM hexamer). All MCMs tested exhibited at least a small degree of substrate unwinding under the assay conditions (Figure 2b; Supplementary Figure 5). Of the non-synthetic enzymes, the two most active MCMs at 25 °C were from mesophilic organisms (*Mac*MCM and *Mba*MCM). *Sso_N_Pfu_C_*MCM showed a similar degree of unwinding to *Mba*MCM, but the synthetic construct lacks a regulatory WHD, which has been shown to have a negative effect on unwinding rate in MCMs^17,18^.

**Figure 2.**
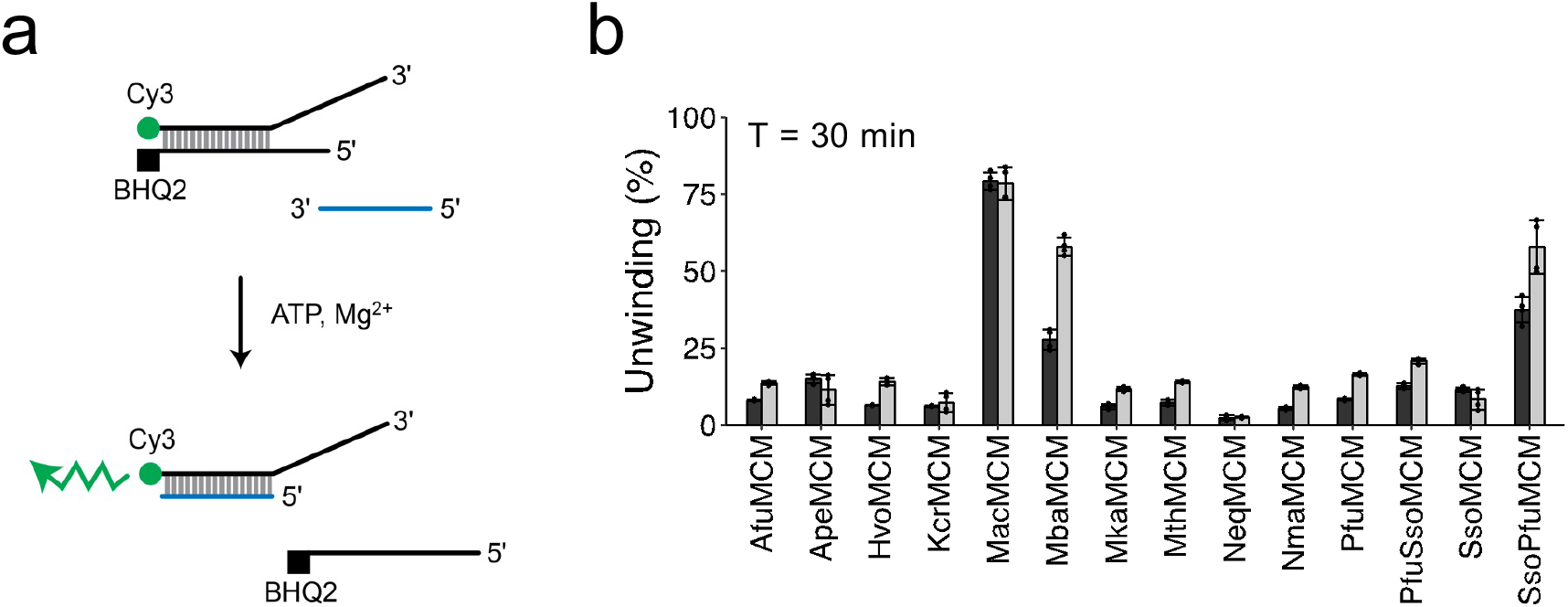
Characterization of DNA unwinding of 14 archaeal MCM helicases. **a** Overview of the FRET-based biochemical activity screen^16^. Protein is pre-equilibrated with a forked DNA substrate. Addition of ATP/Mg^2+^ initiates substrate unwinding, which spatially separates a fluorophore (Cy3) and quencher (BHQ2) causing an increase in fluorescence (λ = 570 nm). A scavenger strand (blue) prevents reannealing. **b** Percentage DNA unwound by each MCM sample at 25 °C (black) and 45 °C (grey) after 30 minutes. Unwinding was quantified by subtracting a no helicase control and then standardizing against a maximum fluorescence well, containing non-annealed Cy3-labelled ssDNA. Bars represent mean unwinding (n = 4). Error bars correspond to ± 1 sem.

MCMs from thermophilic archaea, including *Mth, Pfu* and *Sso*MCM, showed only low activity at 25 °C. Increasing the assay temperature to 45 °C improved activity for most MCMs tested but, even at the elevated temperature, no enzyme was more active than *Mac*MCM. *Mba*MCM also demonstrated high activity at both 25 and 45 °C, however, as expression yields of *Mba*MCM were considerably lower than *Mac*MCM (6 *vs* 92 mg protein per L of culture), we elected to take forward the latter for further study.

### The homohexameric structure of *Mac*MCM resolved at 2.6 Å

To date, all reported 3D structures of homomeric MCMs have been from thermophilic organisms, which demonstrated low activity at 25 °C in our assay. To address this knowledge gap, crystallisation screens were set up using various *Mac*MCM constructs. Crystals that diffracted to 2.6 Å were obtained using a construct that lacked the C-terminal winged-helix domain and carried a point mutation in the Walker B motif (E391Q) to render it inactive (Supplementary Figures 6,7). Crystals were grown in the presence of ATP and MgCl_2_ over a period of 3 days.

The asymmetric unit contained a single ring-shaped *Mac*MCM homohexamer (Supplementary Table 3)^19^. Overall, the structural organisation of *Mac*MCM was consistent with previously published MCM helicases, where each monomer is composed of two modular domains: An N-terminal DNA-binding domain (NTD) and a C-terminal ATPase domain (AAA+) (Figure 3). The NTD is further divisible into 3 subdomains, which consist of a four helix bundle (sA), a four cysteine (C_4_)-type zinc finger (ZnF), and an oligonucleotide binding fold (OB-fold). The active sites of the ATPase domain are formed at subunit-subunit interfaces with both subunits contributing residues, as is typical in MCMs^20^. In each ATPase site, the Walker A, B and Sensor-1 motifs are provided by the *cis-*acting subunit, whilst the Arginine Finger and Sensor-2 motifs are provided by the *trans*-acting subunit. The conserved DNA-binding hairpins are found in the central channel with each subunit providing 3 hairpins: the N-terminal β-hairpin, the Helix-2 Insert and the Pre-sensor-1 β-hairpin^21^.

**Figure 3.**
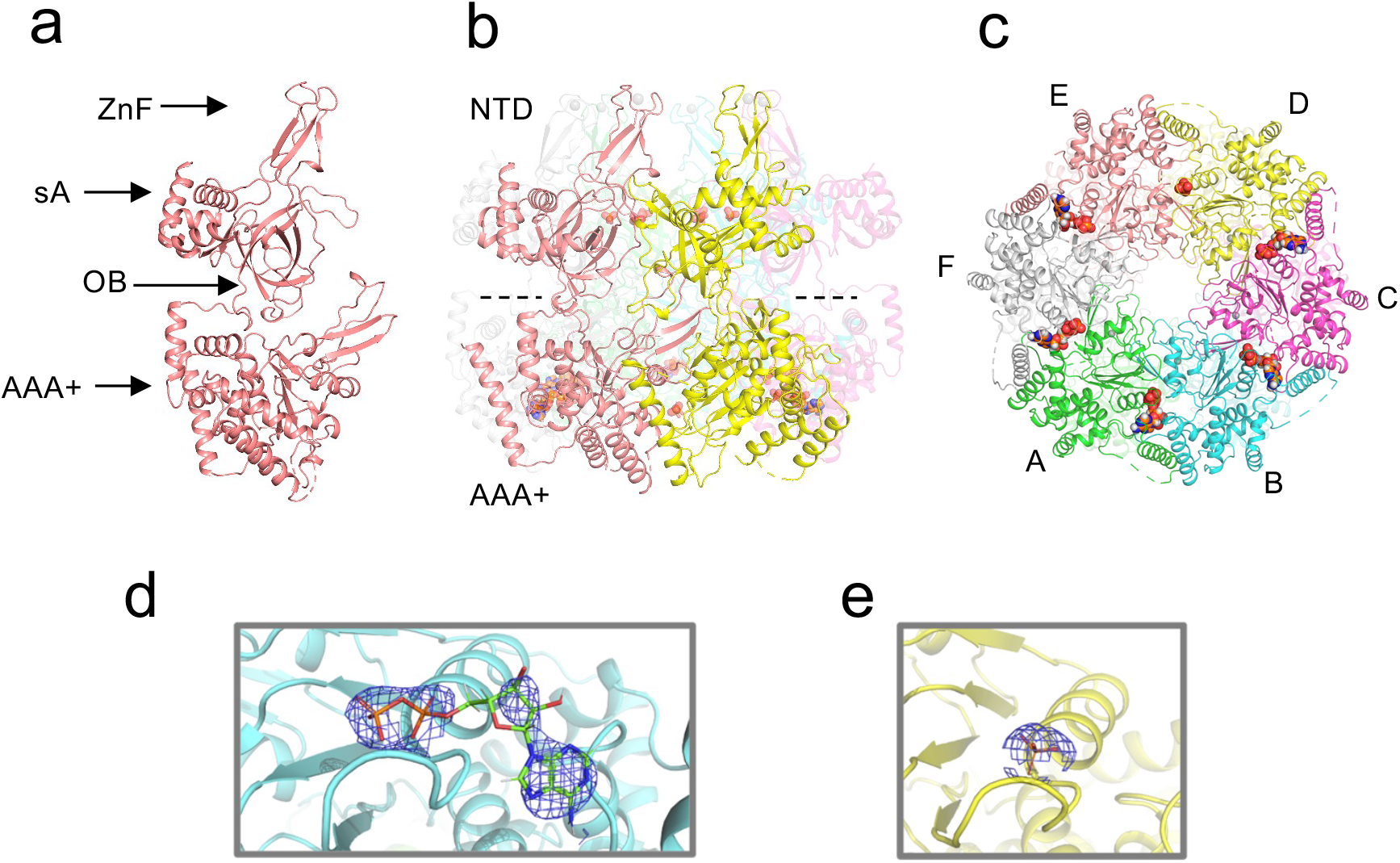
2.6 Å resolution crystal structure of the core *Mac*MCM^ΔWHD E391Q^ hexamer bound to ADP and phosphate. **a** View of a single MCM subunit (chain E), showing the positions of subdomain A (sA), zinc finger (ZnF), oligosaccharide/nucleotide binding fold (OB), and ATPase associated with various cellular activities (AAA+) domain. **b** View perpendicular to the central channel, where the N-terminal domain (NTD) and C-terminal tiers (CTD) are clearly defined. **c** View into the central channel from the CTD side. Position of ADP and phosphate are represented in the sphere format. **d** Close up view of one of the five ADP molecules modelled in an AAA+ active site. ADP is shown in the stick format and electron density (Omit-map) shown in mesh format. **e** Close up view of the phosphate ion modelled in the active site at the interface of subunits D and E. Phosphate is shown in stick format and electron density (Omit-map) shown in mesh format. All images were prepared using PyMol^19^. Each MCM subunit is coloured as stated and represented in the cartoon format.

The structure was solved with four classes of ligand: Zn^2+^ ions were identified in all six ZnF; ADP was identified in five of the six ATPase active sites; a phosphate ion was identified in the remaining ATPase active site (formed by subunits D and E); and six phosphate ions were coordinated by each OB-fold (Figure 3). No density was observed in the ATPase active site for Mg^2+^. Despite the presence of two distinct classes of ligands in the ATPase sites, the structures of the active sites are otherwise indistinct (Supplementary Figure 8), resulting in the DNA-binding hairpins forming a planar orientation with respect to the tiers of the hexamer.

The largest topological difference between *Mac*MCM and other MCM structures concerned the position of the ZnFs. The distance between neighbouring ZnFs in *Mac*MCM is further apart than has been observed in other homomeric MCM hexamer structures (Supplementary Figure 9). While there is evidence of functionally relevant asymmetry in the positioning of ZnFs in MCM2–7, the asymmetry observed in our structure of *Mac*MCM is most likely the result of interactions between molecules in the crystal.

Overall, the 3D structure of *Mac*MCM is consistent with previously published hexameric archaeal and eukaryotic MCMs. When each subunit of *Mac*MCM is superposed in turn with each subunit in a homohexameric MCM (*Sso*MCM, PDB:6MII)^21^ or a MCM2-7 heterohexamer (*Sce*MCM2-7, PDB:6EYC)^22,23^, the average all atom RMSD ± 1 standard deviation is 2.5 ± 0.2 Å and 3.3 ± 0.8 Å, respectively. However, domain specific differences in similarity are observed. The largest differences are found at the N-terminal domain where the structure of *Mac*MCM is more similar to *Sso*MCM (average RMSD: 2.5 ± 0.2 Å) than to *Sce*MCM2-7 subunits (average RMSD: 3.5 ± 1.6 Å). By comparison, the highly conserved C-terminal ATPase domain shares excellent and more consistent structural homology with both *Sso*MCM (average RMSD: 1.3 ± 0.1 Å) and *Sce*MCM2-7 (average RMSD: 1.8 ± 0.2 Å).

### The subunit interfaces of *Mac*MCM are similar to eukaryotic MCMs

The spiral staircase model of DNA translocation by MCMs is dependent on efficient movement of subunits within the hexameric ring^21^. Enzymes from thermophilic organisms typically contain a higher number of salt bridge interactions, which has been proposed to increase stability at higher temperatures^24,25^. However, at lower temperatures, a larger number of salt bridges may rigidify enzymes to an extent that impedes efficient conformational change and thus activity. We analysed subunit interfaces from the hexameric MCM structures of *Sso*MCM (PDB: 6MII)^21^, *Sce*MCM2–7 (PDB:6EYC)^22,23^ and *Mac*MCM (this study) using Proteins Interfaces Structures and Assemblies (PISA)^26^. Despite each eukaryotic *Sce*MCM2-7 subunit having on average 31 % more amino acids than *Sso-* or *Mac*MCM, the oligomerization interface of all three MCMs are formed from equivalent numbers of residues. However, subunit-subunit interfaces in both *Sce*MCM2-7 and *Mac*MCM possess 25% fewer hydrogen bonds and half the number of salt-bridges compared to *Sso*MCM (Supplementary Figure 10). While limited by the availability of equivalent crystal structures for archaeal hexameric MCMs, this analysis is consistent with adaptation to high temperature environments involving an increased number of subunit-subunit interactions^24^. Furthermore, these data suggest that there are greater structural similarities between *Mac*MCM and eukaryotic *Sce*MCM2-7 than between *Mac*MCM and previously studied thermophilic MCMs.

### *Mac*MCM homohexamer unwinds DNA with sigmoidal kinetics

Analysis of the dsDNA unwinding properties of *Mac*MCM revealed a sigmoidal profile, with maximum activity observed after ∼6 minutes when measured at 25 °C (Figure 4a). By contrast, *Sso*_N_*Pfu*_C_MCM exhibited more standard reaction kinetics (Supplementary Figure 11). This behaviour was not observed for any of the other MCMs tested. Sigmoidal enzyme kinetics suggest the presence of a primary, rate-limiting step that precedes DNA unwinding. Such a step has been well described for MCM2-7, where slow ATP-dependent conformational changes in MCM2 and MCM5 occur on a time scale of 5-10 minutes and permit stable assembly of the full hexameric complex on DNA^6,27^. *In vitro*, assembly of the MCM2-7 hexamer is also dependent on protein concentration, ATP binding, temperature and salt concentration^8^. An equivalent behaviour has not been reported for a homohexameric MCM.

**Figure 4.**
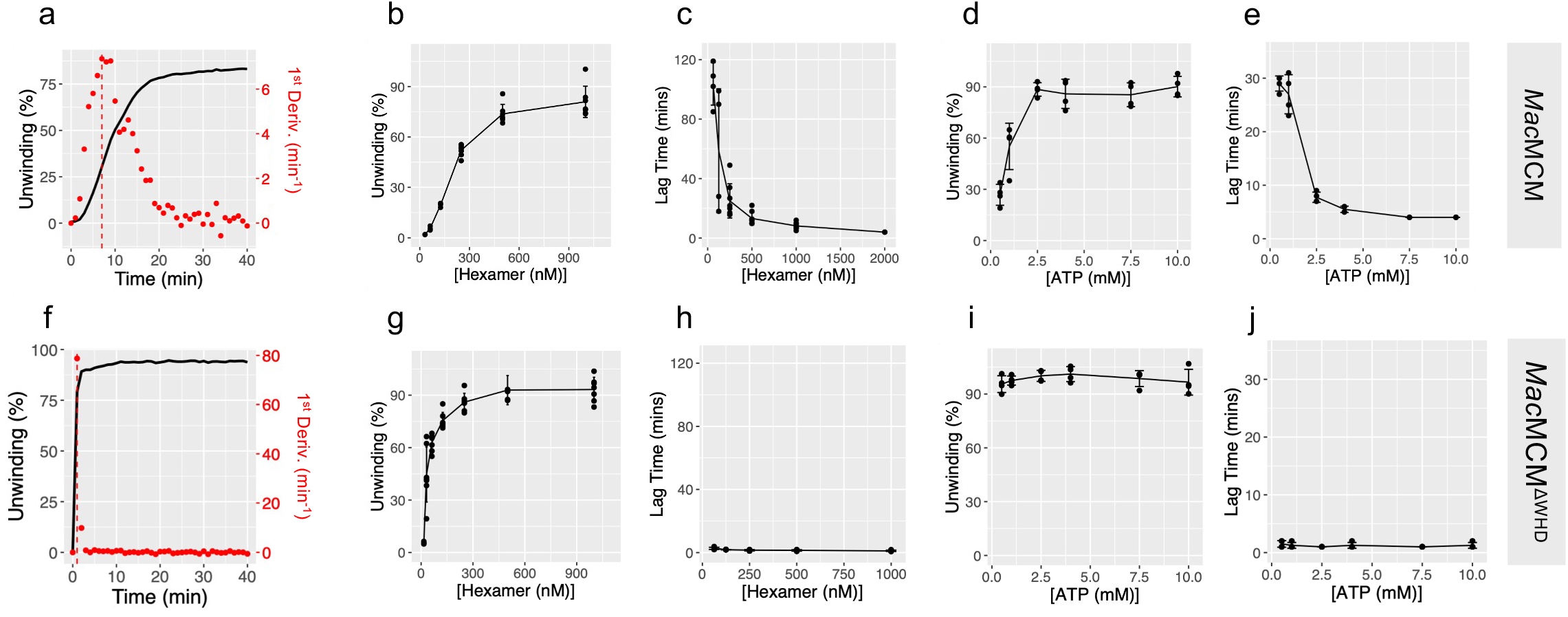
Interactions with protein, ATP and the winged-helix domain influence a slow kinetic step for *Mac*MCM assembly. Assay data and analysis for (a-e) *Mac*MCM and (f-j) *Mac*MCM^ΔWHD^. a,f) Example lag time calculation for a real-time helicase trace for *MacMCM*. The first derivative is calculated from an experimental unwinding curve and the time taken to reach the maximum rate is extracted (red dotted line). b,g) Net unwinding data for *Mac*MCM measured against protein concentration. c,h) Lag time extracted from each protein concentration in part (b). d,i) Net unwinding data for *Mac*MCM measured against ATP concentration. e,j) a “lag time” extracted from each ATP concentration in part (d,i). Concentration when reagent was fixed: MCM hexamer, 1 μM; forked DNA substrate, 50 nM; ATP and MgCl_2_, 4 mM and 10 mM respectively.

To determine which factors influence the activity of *Mac*MCM, DNA unwinding assays were performed using a series of different protein and ATP concentrations. To quantify the sigmoidal kinetics of *Mac*MCM, the first derivative of the measured unwinding curve was calculated and from that the time taken to reach the maximum unwinding rate determined (Figure 4a). We refer to this metric as ‘lag time’.

Decreasing the concentration of *Mac*MCM results in lower relative enzymatic activity and a longer lag time (Figure 4b-c). The effect of ATP on lag time was also measured against a physiological range of ATP concentrations (0.5-10 mM). Reducing the ATP concentration from 2.5 to 0.5 mM markedly decreased net enzyme activity and extended the lag time towards the time limit of the experiment (Figure 4d-e). Together, these results show that both protein-protein and protein-ATP interactions influence the kinetics of *Mac*MCM activity. The observation that higher concentrations of either ATP or protein decrease the lag time suggests that this phenomenon is related to an association event, e.g., the formation of a macromolecular MCM complex.

### Sigmoidal kinetics of *Mac*MCM are influenced by the winged helix domain (WHD)

Cryo-EM studies have demonstrated that the slow ATP-dependent conformational change in MCM2–7 involves repositioning of the WHD of MCM5, which results in stabilisation of the heterohexamer-DNA complex^10,11,28^. In a heterohexamer it is straightforward to assign conformational changes to individual subunits. However, it is unknown whether or how this behaviour evolved from a system in which all subunits are identical. It is also unclear whether all subunits and all subunit-subunit interfaces in a homohexamer behave like the MCM2-MCM5 interaction, or what effect the presence of multiple equivalent WHDs has. To address these issues, we evaluated the DNA unwinding properties of a truncated variant of *Mac*MCM that lacks the WHD (*Mac*MCM^ΔWHD^) across a range of protein and ATP concentrations (Figure 4f-j). Consistent with studies of other MCMs, removal of the WHD resulted in an increase activity with close to 100% of the substrate unwound in the first few minutes (Figure 4f)^17,18^. Moreover, the *Mac*MCM^ΔWHD^ construct did not exhibit a measurable lag time at any of the protein or ATP concentrations tested (Figure 4h,j). These data show that removal of the WHD either eliminates the lag time or substantially reduces it beyond the detection limit of our assay. Nevertheless, a plausible explanation is that the WHDs of *Mac*MCM is directly involved in the rate limiting step in a similar manner to that previously reported for the WHD of MCM5 in the assembly of MCM2-7.

### Both ATP and DNA are required for *Mac*MCM to form a homohexamer

Previously studied archaeal MCMs form higher order homo-oligomers (typically hexamers or dodecamers) even in the absence of ATP or DNA^18,20,29^. In eukaryotes, however, ATP and DNA are required to stabilise a hexameric state of MCM2-7^8^. To evaluate the oligomeric state of *Mac*MCM in the presence of DNA, we incubated the helicase with different ligands before subjecting the samples to analytical size exclusion chromatography (SEC). Single stranded (ssDNA) was used in these experiments to allow us to characterise complexes formed on DNA in the absence of any unwinding.

In the absence of ATP or ssDNA, *Mac*MCM and *Mac*MCM^ΔWHD^ eluted at larger volumes than would be expected for a homohexamer (Figure 5a,b). This was further confirmed using SEC MALLS, which showed that both constructs eluted with molecular weights closer to a monomer, and that the molecular weight calculated was dependent on the concentration of protein loaded (Supplementary Figure 12). This latter observation suggests that apo-*Mac*MCM exists in a mono-oligomer equilibrium in solution and that the monomeric species predominates at lower protein concentrations and in the absence of ligands. These results also suggest that the core region of *Mac*MCM, comprising NTD and AAA+ domains, is not itself able to form a stable homohexamer in solution. This contrasts the *Mac* enzyme with MCMs from the (hyper)thermophilic archaea *Ape*^30^*, Sso*^18^ and *Mth*^31^.

**Figure 5.**
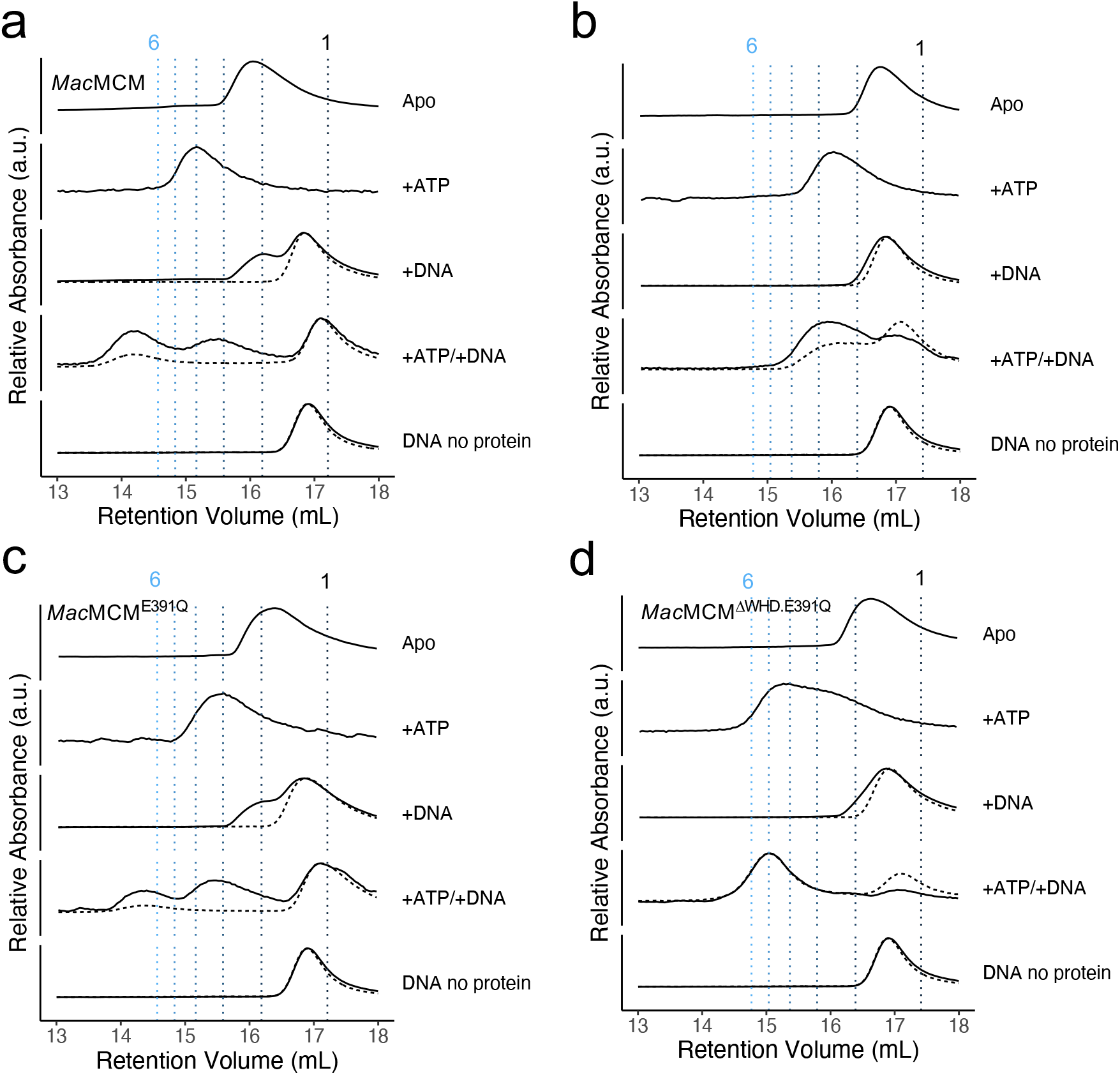
ATP and DNA contribute to stable hexamer formation by *Mac*MCM. The loading of *Mac*MCM constructs on to DNA was analysed by analytical SEC. Protein samples (10 µM) were pre-incubated with or without an equimolar ratio of fluorescein labelled ssDNA substrate (polyT_50_) before application to a Superose 6 Increase 10/300 GL SEC column. Where stated, ATP at 1 mM and Mg^2+^ at 10 mM were added to the buffer. UV absorbance was monitored at both 290 nm (solid trace) and 495 nm (dotted trace). Vertical dotted lines indicate the expected elution volumes of MCM oligomers with 1 to 6 subunits. **a** *Mac*MCM, **b** *Mac*MCM^ΔWHD^, **c** *Mac*MCM^E391Q^ and **d** *Mac*MCM^E391Q.ΔWHD^

The elution profile of *Mac*MCM depends on which ligands are present (Figure 5). The addition of ssDNA does not change the elution volume of either *Mac*MCM or *Mac*MCM^ΔWHD^ or of the ssDNA ligand. In the presence of ATP/Mg^2+^, both *Mac*MCM and *Mac*MCM^ΔWHD^ show a slight decrease in elution volume, but the size of the change does not support the formation of a homohexamer. Only when both ssDNA and ATP are present does the elution profile of *Mac*MCM show a species that elutes at a volume more consistent with a homohexamer (Figure 5a). Moreover, under these conditions *Mac*MCM co-elutes with ssDNA. By contrast, when *Mac*MCM^ΔWHD^ is mixed with ssDNA and ATP the elution volume decreases but not by the same degree as *Mac*MCM. As the ΔWHD variant is considerably more active than the wild-type *Mac*MCM, we hypothesize that higher enzymatic turnover of ATP during the SEC experiment reduces the lifetime of a full *Mac*MCM^ΔWHD^-ATP-ssDNA complex such that a hexameric MCM/DNA species is not resolved. To evaluate the impact of ATP hydrolysis, SEC experiments were repeated using a Walker B mutation (E391Q) that renders the ATPase domain inactive but that should still permit ATP binding^32^. In the presence of ATP but with hydrolysis no longer possible both *Mac*MCM^E391Q^ and *Mac*MCM^ΔWHD.E391Q^ co-eluted with ssDNA at elution volumes consistent with a homohexamer (Figure 5c-d).

### ATP turnover promotes assembly of a DNA-bound *Mac*MCM homohexamer

Hexamerization is a requirement for stable binding of DNA by MCMs^33^. In agreement with previous studies, we found that *Sso_N_Pfu_C_*MCM is an obligate homohexamer at room temperature and pressure (Supplementary Figure 12) and that it binds to forked DNA in the absence of nucleotide co-factors (58.0 ± 9.8 nM; Table 1; Supplementary Figures 13, 14). Adding ATP increases DNA-binding affinity three-fold to 18.3 ± 1.0 nM compared to apo *Sso_N_Pfu_C_*MCM. Incubating *Sso_N_Pfu_C_*MCM with either non-hydrolysable ATP analogues (AMP-PCP, which mimics the pre-hydrolysis state; or ATP-AlF_4_^-^, which mimics the transition state) or ADP did not significantly affect the affinity for DNA.

**Table 1.** DNA binding affinity of MCM constructs is differentially impacted by nucleotides. Binding affinities were measured via fluorescence polarisation using a fluorescently labelled forked DNA substrate in the absence of presence of nucleotides that mimic steps of the catalytic cycle. n.d., not determined. Reactions were conducted using 1 nM DNA and 4 mM nucleotide plus 10 mM MgCl_2_ when added.

| Ligand | E391Q | State | K<sub>d</sub> (nM) |  |  |
| --- | --- | --- | --- | --- | --- |
|  |  |  | *MacMCM* | *MacMCM*<sup>ΔWHD</sup> | *SsoN**PfuC**MCM* |
| - | × | no nucleotide | 476.3 ± 34.5 | > 1000 nM | 58.2 ± 9.4 |
| AMP-PCP | × | pre-hydrolysis | 700.9 ± 70.7 | > 1000 nM | 55.2 ± 3.9 |
| ATP | ✓ | pre-hydrolysis | 780.5 ± 83.7 | 114.3 ± 5.4 | n.d. |
| ADP-AlF<sub>4</sub> | × | transition state | 660.3 ± 62.0 | > 1000 nM | 43.8 ± 6.5 |
| ADP | × | post-hydrolysis | 567.8 ± 60.0 | > 1000 nM | 56.4 ± 2.1 |
| ATP | × | active hydrolysis | 115.0 ± 8.4 | 60.4 ± 2.1 | 18.3 ± 0.9 |

Compared to *Sso_N_Pfu_C_*MCM*, Mac*MCM binds to a forked DNA substrate with more moderate affinity in the absence of ATP (*K*_d_ = 476 ± 34.5 nM). Addition of AMP-PCP, ATP-AlF_4_^-^ or ADP slightly reduced DNA binding affinity compared to apo *Mac*MCM (Supplementary Figure 13). A Walker B mutant of *Mac*MCM, which does not turnover ATP, interacts with DNA with an affinity of 780 nM in the presence of ATP. However, a large increase in affinity for DNA is seen when the enzyme is capable of turning over ATP (*K*_d_ = 115 ± 8.1 nM). This represents a 4-fold change compared to apo-*Mac*MCM and an 8-fold changed to compared to the E391Q variant in the presence of ATP.

Truncation of the WHD of *Mac*MCM decreased affinity for DNA in all conditions tested except for when ATP was present (Table1). Apo-*Mac*MCM^ΔWHD^ bound DNA with much weaker affinity compared to the apo full-length protein (Table 1; Supplementary Figure 13c). Likewise, weak binding to DNA was seen in the presence of AMP-PCP, ATP-AlF_4_^-^ or ADP. However, compared to wild-type *Mac*MCM, a smaller 2-fold difference in affinity for DNA was seen between active (K_d_ = 60.4 ± 2.1 nM) and catalytically inactive (K_d_ = 114.3 ± 5.4 nM) variants of *Mac*MCM^ΔWHD^. In both cases, the affinity of the WHD truncation for DNA is higher than the full-length *Mac*MCM.

## Discussion

All replicative helicases are believed to unwind DNA as ring-shaped hexamers. However, each helicase family functions differently and there is no universally conserved mechanism that is used to load each class of hexamer onto DNA. In bacteria, a closed DnaB homohexamer is recruited to double stranded DNA by the cofactor proteins DnaA and DnaC^34,35^. DnaC is able to break open the closed-form DnaB homohexamer to allow loading onto dsDNA^35^. In eukaryotes, MCM2-7 is recruited to DNA as an open heterohexamer. The open form of MCM2-7 is stabilised through interaction with Cdt1, which is then recruited to DNA by ORC and Cdc6^8,10,11,28^. By contrast, the mechanism by which homohexameric archaeal MCM helicases load onto dsDNA has so far proven elusive. The lack of an experimentally tractable system has prevented a comparison of homomeric vs heteromeric MCMs, as well as limiting a broader understanding of the properties of replicative helicases across the three domains of life.

Archaeal MCMs are predominantly thought to form homohexamers and, while being simpler in composition than the heteromeric MCM2-7 complex, a homomeric composition poses fundamental questions about assembly and which aspects of MCM biology are conserved. Despite considerable recent advances in our understanding of the assembly of the eukaryotic MCM2-7 complex, we still have few details concerning how archaeal homohexamer MCMs assemble on DNA and what similarities and differences exist between the eukaryotic and archaeal systems. Archaea possess homologues of the eukaryotic MCM recruitment factors ORC and Cdc6, but they do not universally possess homologues of cofactors that function to break a ring (DnaC) or stablise an open conformation (Cdt1). Our appreciation of biochemistry and structural biology of archaeal MCMs is largely limited to the properties of proteins from two thermophilic organisms, *M. thermautotrophicus* and *S. solfataricus*^36^.

In the 25 years since archaeal MCM were first characterised, new archaea lineages have been identified that inhabit diverse environments and with evermore closer evolutionary links to eukaryotes^37^. Here, we reasoned that characterising the activity of MCMs across a broader range of archaeal organisms may identify candidates that provide a better model for investigating the assembly of a homomeric MCMs under ambient experimental conditions. Our current work provides such a system – *Mac*MCM, the sole MCM from the ectosymbiotic archaea *M. acidophilum*. Investigating *Mac*MCM has allowed us to gain clearer insight into the mechanisms that permit assembly of a homomeric MCM onto a synthetic DNA substrate.

### *Mac*MCM shows similar properties to eukaryotic MCMs

Like other archaeal MCMs, *Mac*MCM can unwind dsDNA in the absence of protein co-factors. However, real-time analysis of the unwinding dynamics of *Mac*MCM revealed multiphase kinetics in which a slow initial phase precedes a rapid second phase (Figure 4a). *Mac*MCM therefore shows strong parallels with the rate-limiting kinetics of the closure of the eukaryotic MCM2-5 gate, which precedes the formation of a stable, DNA-bound MCM2-7 heterohexamer (Figure 6)^6^. In particular: (1) Both events occur on a comparable time scale (∼5-10 minutes), which cannot be attributed to diffusion alone ^6^; (2) Apo *Mac*MCM shows weak self-association (Figure 5a), while MCM2 and 5 subunits do not co-elute *in vitro* ^38^; (3) ATP binding is required to stabilise association of subunits in *Mac*MCM (Figure 5a) and MCM2-7 ^8^; (4) Both the kinetics and formation of a hexamer and its stability are influenced by protein or ATP concentration (Figure 4)^6,8^;(5) Efficient assembly of *Mac*MCM on a forked DNA substrate requires ATP hydrolysis rather than just ATP binding (Figure 5; Table 1)^6,11^; and (6) the transition from open to DNA-bound closed complex is dependent on the WHD (Figure 4)^11,28^. It is important to note that like our study, many of these observations for MCM2-7 were also made in absence of co-factors^6^ on a range of different DNA substrate topologies. Observation of these critical assembly details for a homohexameric MCM has only been possible because we were able to identify an MCM for which the unliganded state shows monomer-like properties.

**Figure 6.**
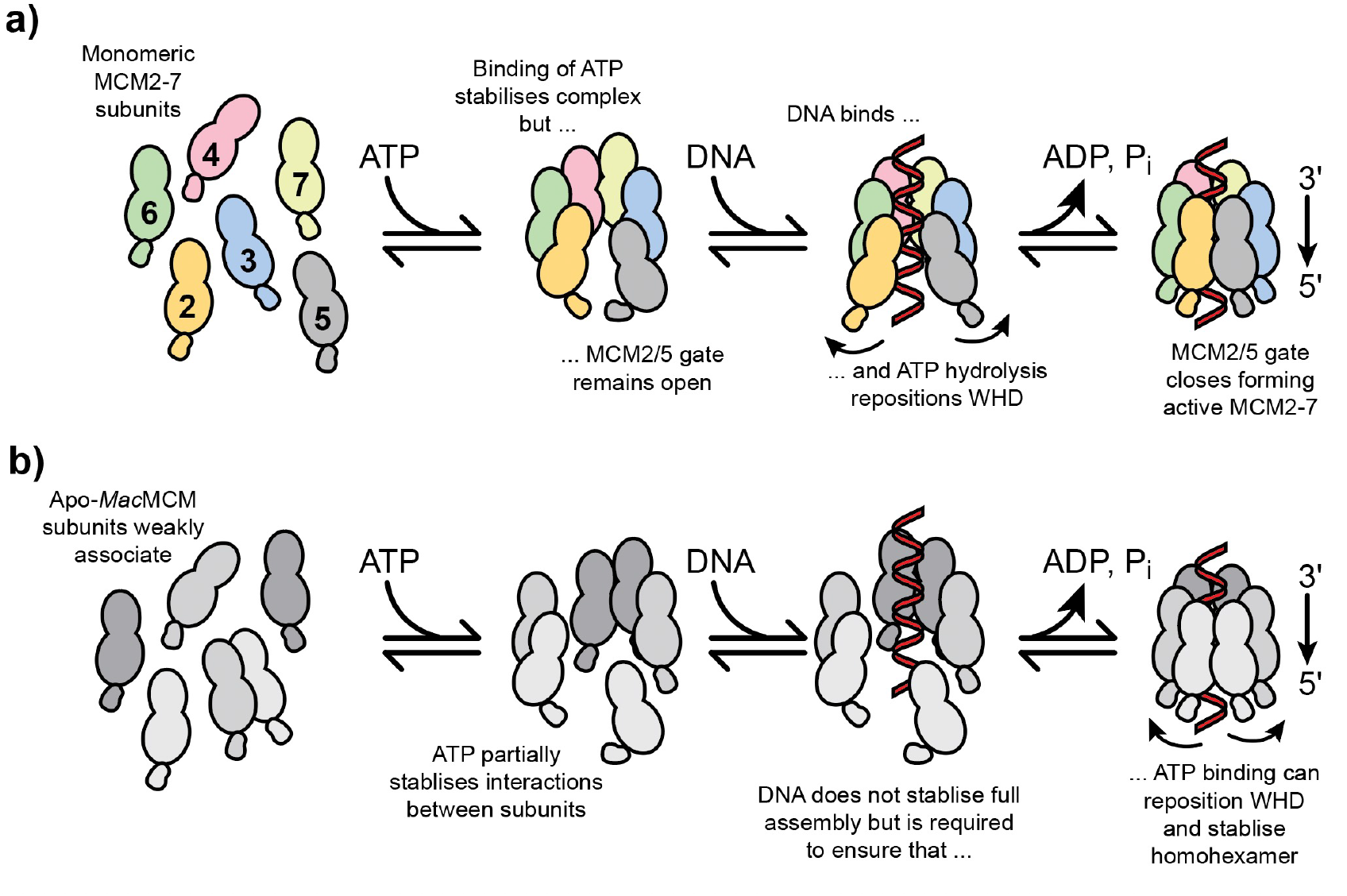
Conservation of core steps in the assembly of homo- and hexameric MCM complexes. **a** Binding of ATP stabilises interactions of MCM2-7 subunits. Open-form MCM2-7 interacts transiently with ssDNA until ATP hydrolysis repositions the WHD of MCM5 to allows closure of the MCM2-5 gate. **b** *Mac*MCM subunits only weakly associate in the absence of ligands. Binding of ATP alone only partially stabilises the interaction between *Mac*MCM subunits, but not sufficiently to form stable homohexamers. ATP binding is required for the formation of a homohexameric complex with DNA. Hydrolysis of ATP is not necessary as an inactive enzyme (E391Q mutant) is able to interact with DNA. ATP binding is potentially required to to reposition one or more of the WHDs.

The affinity of MCMs for DNA is impacted by the presence of nucleotides and the WHD. In MCM2-7, ATP hydrolysis is needed to allow efficient repositioning of the WHD of MCM5 and the subsequent formation of an active MCM2-7 hexamer^11^. The affinity of *Sso_N_Pfu_C_*MCM for DNA increases two-fold in the presence of ATP compared to no nucleotide despite this construct lacking a WHD (Table 1). Apo-*Mac*MCM binds DNA less strongly than the *Sso-Pfu* chimera, however ATP has a stronger effect on affinity. Indeed, constructs of *Mac*MCM that can hydrolyse ATP show the highest affinity for a forked DNA substrate. As seen with MCM2-7, these data suggest that hydrolysis of ATP rather than ATP binding alone appears to be most important for higher affinity interaction with DNA. Moreover, a WHD truncation of *Mac*MCM interacts with forked DNA more strongly than the full-length enzyme in the presence of ATP. These data suggest that, like eukaryotic heteromeric MCMs, homomeric archaeal MCMs also require ATP-driven reposition of the WHD to achieve high affinity binding of DNA.

The genome of *M. acidophilum* contains genes that are homologous to important MCM2-7 loading factors in eukaryotes, including ORC, GINS and RecJ/Cdc45^39^. We did not explore how these co-factors contribute to the assembly of *Mac*MCM or its interaction with DNA substrates. Future characterisation of these co-factors would address important questions around *Mac*MCM function, including its assembly onto a closed-circular genomic DNA *in vivo* as well as elaborating on the similarities between the roles and evolution of co-factors in archaea and eukaryotes.

### Can *Mac*MCM form higher order oligomers?

The formation of an MCM double hexamer is thought to be an important step in replication initiation in both archaea and eukaryotes ^29,40^. We saw no evidence that *Mac*MCM forms a double hexamer in either SEC or SEC MALLS data under any of the conditions tested. Moreover, EMSAs only showed a single mobility shift across the concentration range tested, consistent with a single type of *Mac*MCM-DNA complex. However, the low intrinsic affinity for DNA in the absence of ATP does not preclude the formation of higher order complexes at elevated protein concentrations. Indeed, *Sso*_N_*Pfu*_C_MCM, which bound forked DNA with much higher affinity than *Mac*MCM in our EMSA assays, did show evidence of forming two higher order oligomeric species, presumably a hexamer and double hexamer, even though only a homohexamer was observed by SEC MALLS at similar concentrations. While we do see crystal-induced asymmetry in the positioning of the six ZnFs, the structure and sequence of the N-terminal domain of *Mac*MCM otherwise appear compatible for the formation of a double hexamer similar to those previously described for other archaea and eukaryotic MCMs.

### Are thermophilic proteins good models for mesophilic MCM helicases?

Organisms are uniquely adapted to their environment. This extends to the protein level, where enzymes are adapted to work efficiently in conditions specific to the environmental niche of the organism. Under the ambient conditions of our activity screen, proteins from organisms that inhabit low temperature environments were the most active. Previous studies have demonstrated that *Mth*MCM and *Sso*MCM display high activity at temperatures >50 °C ^21,41^. However, in our assays neither of these enzymes demonstrated substantial activity at temperatures below 45 °C.

The effect of temperature on oligomeric state and stability has been previously explored for both eukaryotic and archaeal MCMs. It was initially found that *Mth*MCM predominantly exists as a dodecamer at room temperature^29^, however when SEC analyses were later performed at higher, near-physiological temperatures (50 °C), *Mth*MCM populations redistributed into hexameric subspecies^42^. Open-form hexamers of both *Mth*MCM and *Sso*MCM were observed by negative stain EM when samples had been pre-heated to temperatures closer to physiological conditions of the organism^43,44^. Similarly, unliganded eukaryotic *Sce*MCM2-7 hexamers dissociate when the temperature is increased from 4 °C to 30 °C^8^.

We determined that the unliganded conformation of *Mac*MCM to be predominantly monomeric in solution, which is the first time such a species has been observed for an active archaeal MCM. These data support the conclusion that oligomeric proteins that are adapted for higher temperature environments show increased assembly stability at ambient temperatures. Indeed, we observe that subunit-subunit interfaces of MCM complexes from thermophilic organisms contain higher numbers of hydrogen bonds and salt bridges than mesophilic counterparts: The interfaces observed in our *Mac*MCM crystal structure are more similar to yeast MCM in terms of polar non-covalent interactions than to an MCM from a thermophilic archaeon. Structure guided mutagenesis on both *Mth* and *Sso*MCM is consistent with these observations. For example, dodecamerisation of *Mth*MCM can be inhibited by mutation of salt bridges that mediate head-to-head hexamer interactions^45,46^. Furthermore, removal of inter-subunit salt bridges in the C-terminal domain of *Sso*MCM, prevents hexamer formation of an unliganded enzyme^20^. Reducing the number of inter-subunit salt bridges in *Sso*MCM also resulted in an increase in helicase activity^20^. It is likely that monomeric species of *Sso*MCM and *Mth*MCM would be detectable if these systems could be studied under conditions closer to those of the environmental niche of these organisms. However, *Mac*MCM provides access to these states under ambient conditions, which offers considerable experimental convenience.

In summary, we report the discovery of an experimentally tractable mesophilic MCM that allows analysis of the assembly pathway of a homohexameric helicase onto a DNA substrate. All extant MCMs have evolved from a homohexameric ancestor but the choice of previous archaeal subjects used in structure-function studies has limited our capacity to probe the differences and similarities that exist between eukaryotic and archaeal MCMs. The data we present here shows that core steps of the assembly pathway are conserved between homomeric and heteromeric MCMs. The fact that similarities exist suggests that the fundamental steps of MCM assembly evolved before the addition of regulatory factors. The structure-function relationship and assembly properties of *Mac*MCM and indeed MCMs from other non-thermophilic archaea thus represent new and important tools for probing the evolution of this critical class of replicative helicase and DNA replication more broadly.

## Methods

### Preparation of recombinant MCM samples

MCM genes were synthesised and cloned into the ampicillin resistant pONT vector by GenScript (Supplementary Table 1). Genes were positioned downstream of a T7 promoter and N-terminally His-10 tagged. All genes were codon optimised for expression in *Escherichia coli* and a double stop codon was added. Where performed, site directed mutagenesis was carried out using QuikChange Lightning Mutagenesis (Agilent), according to the manufacturer’s guidelines. MCM protein was overproduced in *E. coli* BL21 (DE3) pLysS cells (Agilent). Cultures were grown at 37 °C in LB containing 34 μg/mL chloramphenicol, 100 μg/mL ampicillin, 1% (w/v) glucose. When cultures reached an OD_600_ of 0.6-0.8, expression was induced with 1 mM IPTG and placed at 20°C. After 20 hours, the final OD_600_ was measured, and cells were harvested via centrifugation at 4,000 × *g*. Cell pellets were then stored at -80°C until required.

### Purification of Recombinant MCMs

Cell pellets (from 100 mL culture) were thawed and re-suspended in buffer A (20 mM Tris-HCl pH 8.0, 500 mM NaCl, 20 mM imidazole, 5 % (w/v) glycerol) to an OD_600_ of 100. Buffer A was supplemented with DNase, RNase (both at 20 µg/mL) and cOmplete protease inhibitor tablets (Roche). Cells were lysed by sonication at 70 W (3 s on, 7 s off for 1 min per 100 mL culture). Cell extract was then centrifuged for 45 minutes at 30,000 × *g*, 4 °C. The resulting supernatant was loaded onto a 1 mL HisTrap FF column (GE Healthcare), pre-equilibrated in Buffer A. The column was then washed with a high concentration salt wash (20 mM Tris-HCl pH 8.0, 2 M NaCl, 20 mM imidazole, 5 % (w/v) glycerol) before being re-equilibrated into Buffer A. Bound protein was eluted from the column with Buffer B (20 mM Tris-HCl pH 8.0, 500 mM NaCl, 500 mM imidazole, 5 % glycerol). At this point, if samples were being used in the initial characterisation screen, fractions were pooled and dialysed against Buffer C (20 mM Tris-HCl pH 8.0, 500 mM NaCl, 5 % (w/v) glycerol) overnight at 4°C. Dialysed protein was spin-concentrated in a Vivaspin 6 MWCO 50,000 (Sartorius) to the desired concentration, then snap frozen in aliquots and stored at -80 °C. For all other samples, positive fractions from Elution B were pooled and TEV protease was added to a ratio of 1 mg TEV: 50 mg His-tagged protein. Fractions were then dialysed against Buffer D (20 mM Tris-HCl pH 8.0, 500 mM NaCl, 1 mM DTT, 5 % glycerol) overnight at 4 °C. Tag cleavage was confirmed by SDS PAGE. Dialysate was then loaded onto a HisTrap FF column and the recombinant protein collected from the flow through. The flow through was then concentrated in an Amicon Ultra-15 50,000 MWCO spin concentrator to ∼10-20 mg/mL and loaded onto a HiPrep 26/60 S200 Size Exclusion Column (GE Healthcare), equilibrated in Buffer D. Fractions were collected and spin concentrated as before to a final concentration ∼7-20 mg/mL. Samples were either snap frozen in liquid nitrogen and stored at -80 °C, or used immediately to set up protein crystallisation experiments.

### Fluorescent Helicase Assay

Helicase unwinding reactions were carried out on a forked DNA substrate, which was formed by annealing a 5ʹ-Cy3-labelled oligonucleotide 5ʹ-**[Cy3]**GGGACGCGTCGGCCTGGCACGTCGGCCGCTGCGGCCAGGCACCCGATGGC(GTTT)_6_-3ʹ; Merck), to a 3ʹ-BHQ2-labelled oligonucleotide (5ʹ-(TTTG)_8_CCGACGTGCCAGGCCGACGCGTCCC**[BHQ2]**-3ʹ; Eurofins). A scavenger oligonucleotide, (5ʹ -GGGACGCGTCGGCCTGGC-3ʹ; Merck) complementary to the duplex region of the BHQ2-labelled strand was added to the reaction in 10-fold excess to prevent reannealing of unwound substrate. Standard reactions containing 1000 nM helicase (based on hexamer molecular weight), 50 nM forked DNA and 500 nM scavenger were monitored at 25 °C for 30 minutes with a sampling frequency of 1 reading per well per minute. Unless stated, the reaction buffer contained 250 mM potassium glutamate, 20 mM potassium phosphate pH 8.0, 1 % glycerol, 4 mM ATP and 10 mM MgCl_2_).

### Crystallization and data collection

Samples of *Mac*MCM^ΔWHD.E391Q^ were dialysed overnight at room temperature into 100 mM NaCl, 20 mM Tris pH 8.0, 0.5 mM TCEP and 5% glycerol. 10 mM ATP, 10 mM MgCl_2_ was added 10 minutes before setting up the crystallization condition. Long plate-shaped crystals grew over 3 days at 20 °C in a sitting drop containing 10 µL of protein solution and 10 µL of well solution (0.03 M NPS, 0.1 M MOPS/HEPES pH 7.5, 10 % w/v PEG 20,000, 20 % v/v PEG MME 550). Crystals were harvested using a cryo-loop (Crystal Cap HP) and flash frozen in liquid nitrogen. Data were collected at Diamond Lightsource Beamline i03 at a wavelength of 0.9763 Å and temperature of 100 K. Data were scaled and integrated using Xia2–DIALS software package to 2.59 Å resolution ^47^.

### Structure solution and refinement

Initial phases were calculated using Phaser molecular replacement software^48^, which placed six copies of a no loop, polyalanine model of the AAA+ domain of *SsoPfu*MCM in a ring (PDB: 4R7Y)^15^. Following the placement of this model, the electron density map was improved using RESOLVE density modification^49^. Loops and the entire N–terminal domain were then built iteratively using manual building in Coot and the automated software Autobuild and Buccaneer^50–52^. Refinement was carried out using phenix.refine^53^. Following the building of the N-terminal domain, zinc ions were placed through observation of weak anomalous data. Cysteine co-ordination and ligand restraints were then generated using ReadySet!^54^. Where present, nucleotide in the active sites was modelled as ADP.

### Analytical Size Exclusion Chromatography

A Superose 6 Increase 10/300 GL was pre-equilibrated with 200 mM NaCl, 20 mM Tris pH 8.0, 5 % glycerol on an ÄKTA Pure (GE Life Sciences). Where nucleotide was present, the running buffer also included 1 mM ATP and 10 mM MgCl_2_. MCM was diluted to 60 µM monomer concentration in running buffer. Where ligands were present, 5 mM ATP, 10 mM MgCl_2_ or 10 µM DNA was added to the sample. 100 µL samples were loaded and run at 0.5 mL/min. Absorbance was simultaneously measured at 290 and 495 nm. The column was calibrated using thryglobulin (660 kDa), β-amylase (223 kDa), alcohol dehydrogenase (150 kDa), carbonic anhydrase (30 kDa) and cytochrome C (12 kDa; all from Merck). The relationship between elution volume and molecular mass determined using linear regression. Fluorescein-labelled ssDNA substrate (5’[FAM]-pT_50_; Merck) was used in all SEC experiments to allow us to characterise complexes formed on DNA in the absence of any unwinding.

### Size Exclusion Chromatography with Multi Angle Laser Light Scattering (SEC-MALLS)

A Superose S6 Increase 10/300 GL analytical column (GE Healthcare) was equilibrated overnight with 200 mM KCl, 50 mM Tris-Cl pH 8.0, 5 % (*v/v*) Glycerol, 0.5 mM DTT buffer on a Shimadzu HPLC system. A total of 100 µL protein at 1-10 mg/mL was passed over the column at a flow rate of 0.5 mL/min. Light scattering was determined using a Wyatt HELEOS-II multi angle light scattering detector. DRI was determined using a Wyatt rEX refractive index detector. Data were analysed using Astra 7 (Wyatt) software, where the MW is calculated from a Zimm model. BSA run at 2.5 mg/mL was used to normalize the DRI signal. The *dn/dc* value was adjusted until the expected MW of BSA (66 kDa) was obtained.

### Fluorescent anisotropy

Helicase and DNA mixes were prepared in 250 mM potassium glutamate, 20 mM Tris pH 8.0. A forked DNA substrate was prepared by annealing a 5ʹ-FAM labelled oligonucleotide (**5ʹ-[FAM]**GGGACGCGTCGGCCTGGCACGTCGGCCGCTGCGGCCAGGCACCCGATGGC(GTTT)_6_-3ʹ; Merck), with a partially complementary oligonucleotide (5ʹ-(TTTG)_8_CCGACGTGCCAGGCCGACGCGTCCC-3ʹ; Merck). Forked DNA was added to 1 in 2 serially diluted MCM samples with each well containing a final concentration of 1 nM DNA. Where present, binding reactions were supplemented with 4 mM nucleotide and 10 mM MgCl_2_.

### Electrophoretic mobility shift assays

0.8 % (*w/v*) agarose gels were prepared in 1 x Tris-Borate (90 mM Tris, 90 mM Borate, pH 8.3) buffer. Protein and buffer were prepared in 250 mM KGlu, 20 mM Tris-Cl pH 8.0. A forked DNA substrate was prepared by annealing a 5ʹ-FAM labelled oligonucleotide (**5ʹ-[FAM]**GGGACGCGTCGGCCTGGCACGTCGGCCGCTGCGGCCAGGCACCCGATGGC(GTTT)_6_-3ʹ; Merck), with a partially complementary oligonucleotide (5ʹ-(TTTG)_8_CCGACGTGCCAGGCCGACGCGTCCC-3ʹ; Merck). Forked DNA was added to 1 in 2 serially diluted MCM samples with each sample containing a final concentration of 10 nM DNA. A DNA-only control was also prepared to determine the motility of unbound DNA. Samples were then incubated at room temperature for 30 minutes. Before loading, 20 µL 2 x TB and 25 % (v/v) glycerol was added to each sample. 10 µL each sample was then run on the agarose gel for 20 minutes at 150 V. Gels were imaged on a Typhoon scanner (GE Healthcare) using Cy2 filters with a 100 µm imaging pixel size. To estimate the equilibrium dissociation constant (K_d_) between protein and DNA, the protein concentration is identified where the DNA motility is distributed equally between free and protein-bound states.

### NMR spectroscopy

NMR spectroscopy was used to confirm ATPase activity of purified MCM samples. Each reaction mixture (600 µL) contained 50 mM ATP, 2.5 mM MgCl_2_, 10 % (v/v) D_2_O, 250 mM KGlu and 10 mM Tris-Cl pH 8.0. Reactions were initiated by adding MCM to a final concentration of 50 µM (monomeric). After 30 minutes at 25 °C, 1D [^31^P]-NMR spectra were recorded on a 500 MHz NMR spectrometer (Bruker). Spectra were processed using TopSpin software (Bruker).

## Data availability

The atomic model described in this study and accompanying structure factors have been deposited to the Protein Data Bank under the accession code 8Q67.

## Acknowledgements

We acknowledge funding from BioProNET, a BBSRC NIBB co-sponsored by the EPSRC (BB/L013770/1), BBSRC Impact Acceleration Award (BB/S506795/1), Oxford Nanopore Technologies (ONT), and the Department of Biology, University of York. ON was a recipient of a PhD studentship the BBSRC White Rose Doctoral Training Programme (BB/M015831/1). We thank Dr Andrew Leech for assistance and support with biophysical experiments; Dr Alex Heyam for support with NMR spectroscopy; Dr Johan Turkenburg and Sam Hart for assistance with X-ray crystallography; staff at ONT for helpful discussion; Dr Cyril Sanders and Dr Michelle Hawkins for critical reading of the manuscript; and the many undergraduate students who have worked on various aspects of this project over the past 10 years.

