## Supplementary Figures for "Fundamental steps in Minichromosome Maintenance complex assembly are conserved from archaea to eukaryotes"

### Supplementary Material

Supplementary Figures 1-13

Supplementary Tables 1-3

**Supplementary Table 1: The optimal growth temperatures of archaeal species targeted in this study.**

* Optimal growth temperature data is limited for the organism. The conditions of the habitat where the organism was initially identified is taken as a proxy for the organisms potential growth range.

† Full gene contains a highly conserved internal intein that is removed in our construct.

| | | | | | Temperature (ºC) | | | | |
|---|---|---|---|---|---|---|---|---|---|
| Organism | MCM | UniProt Accession Code | $N_{res}$ | MW | Min | Optimum | Max | Phyla | Reference |
| *Archaeoglobus fulgidus* | *Afu*MCM | Q7ZAA5 | 698 | 78,784 | 60 | 83 | 85 | Euryarchaeota | [1] |
| *Aeropyrum pernix* | *Ape*MCM | Q9YFR1 | 697 | 78,472 | 70 | 90 | 97 | TACK | [2] |
| *Haloferax volcanii* | *Hvo*MCM | D4GZG5 | 702 | 78,855 | 30 | 42 | 55 | Euryarchaeota | [3] |
| *Korarchaeum cryptofilum* | *Kcr*MCM | A2BL91 | 703 | 79,373 | 55 | 85 | 90 | TACK | [4] |
| *Mancarchaeum acidophilum* | *Mac*MCM | A0A218NN99 | 687 | 76,126 | 10 | 37 | 45 | DPANN | [5] |
| *Methanosarcina barkeri* | *Mba*MCM | A0A0E3QYF9 | 700 | 78,909 | 30 | 35 | 45 | Euryarchaeota | [6] |
| *Methanohalophilus halophilus* | *Mha*MCM | A0A1L3PZK3 | 696 | 78,154 | 30 | 40 | 55 | Euryarchaeota | [7] |
| *Methanopyrus kandleri* | *Mka*MCM | Q8TWR7 | 656 | 74,062 | 84 | 98 | 110 | Euryarchaeota | [8] |
| *Methanothermobacter thermautotrophicus* | *Mth*MCM | O27798 | 666 | 75,553 | 40 | 65 | 75 | Euryarchaeota | [9] |
| *Nanohaloarchaea archaeon SG9* | *Nac*MCM | A0A1D8MR82 | 675 | 76,073 | 16 | 33 | 40 | DPANN | [*] |
| *Nanoarchaeum equitans* | *Neq*MCM | Q74MT7 | 657 | 74,097 | 75 | 80 | 95 | DPANN | [10] |
| *Nitrosopumilus maritimus* | *Nma*MCM | A9A310 | 695 | 78,136 | 15 | 28 | 32 | TACK | [11] |
| *Pyrococcus furiosus* | *Pfu*MCM | Q8U3I4 † | 681 | 76,704 | 70 | 95 | 103 | Euryarchaeota | [12] |
| *Saccharolobus solfataricus* | *Sso*MCM | Q9UXG1 | 686 | 77,428 | 55 | 75 | 90 | TACK | [13] |

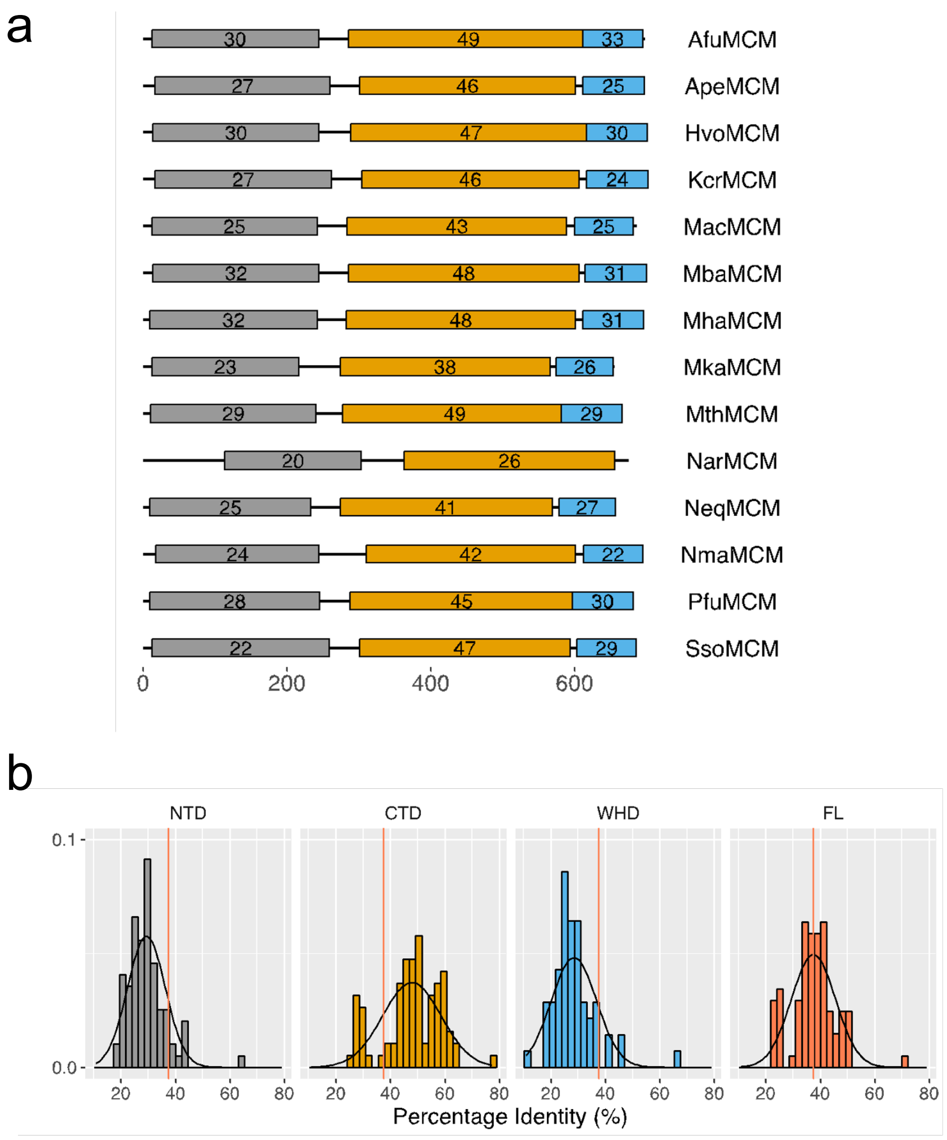


**Supplementary Figure 1: Schematic and conservation analysis of MCM used in this study.** **a** Conserved MCM subdomains domains were identified using the InterProScan[14] tool (https://www.ebi.ac.uk/interpro/search/sequence/). Coloured boxes indicate the identified domain and length (amino acids), where grey is a N-terminal DNA binding domain, orange is a P-type NTPase and blue is a winged-helix domain (WHD). Numbers inside the box indicate the average percentage identity score for each MCM subdomain when aligned to all MCMs studied here (https://www.ebi.ac.uk/Tools/msa/clustalo/). **b** The distribution of percentage identity scores for all MCM and subdomains. The black line represents the normal distribution, with a standard deviation σ and mean μ. The red line represents the mean value for the full-length enzyme. **NTD**: μ=29.4, σ=6.9; **CTD**: μ=47.9, σ=10.7; **WHD**: μ=28.5, σ=8.3; **FL**: μ=37.5, σ=8.1).

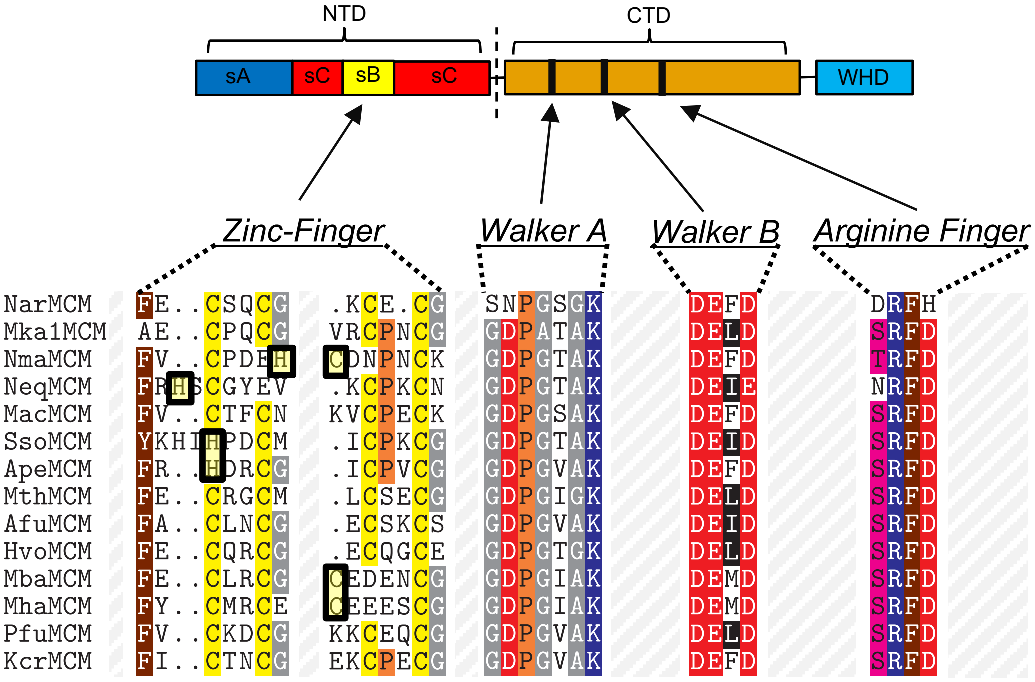


**Supplementary Figure 2: Sequence alignment of core MCM motifs.** MSA of MCM used in this study. Sequences were aligned using Clustal Omega[15] and visualized using the TexShade package in LaTeX. Conserved amino acids are coloured based on the chemical properties of the functional side chain group. The conserved glutamate (E) residue in the Walker B motif is mutated to glutamine (Q) for *Mac*MCM$^{E391Q}$. Brown: aromatic, Yellow: sulphur, Orange: imino, Grey: aliphatic (small), Red: acidic, Blue: basic, Black: aliphatic, Purple: hydroxyl.

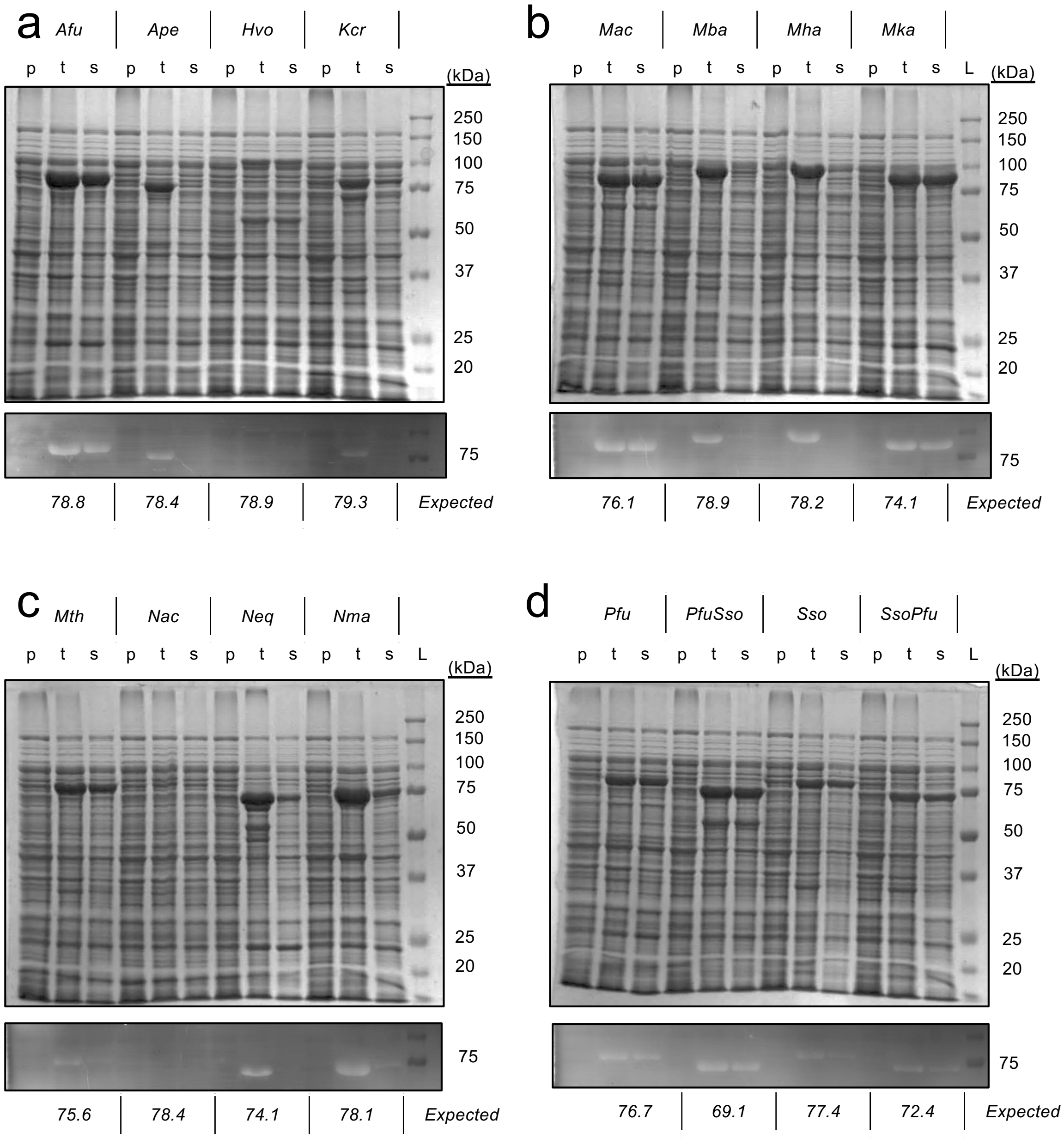


**Supplementary Figure 3.: Test for expression and solubility of 16 archaeal MCM orthologues. a-d** All samples were analysed on a 12 % (*w/v*) SDS-PAGE. Samples represent pre-induction of expression (p), then total (t) and soluble (s) fractions which are collected 20-hours after the IPTG- induction at 20 °C. Top panels represent gels where proteins are stained through a Coomassie- based approach. Bottom panels represent gels where proteins are stained using a fluorescent Ni-NTA conjugate that indicates the presence of poly-histidine tags. Reference (L) lanes represent MW standard marker (Precision Plus Protein™ All Blue Pre-Stained Protein Standards).

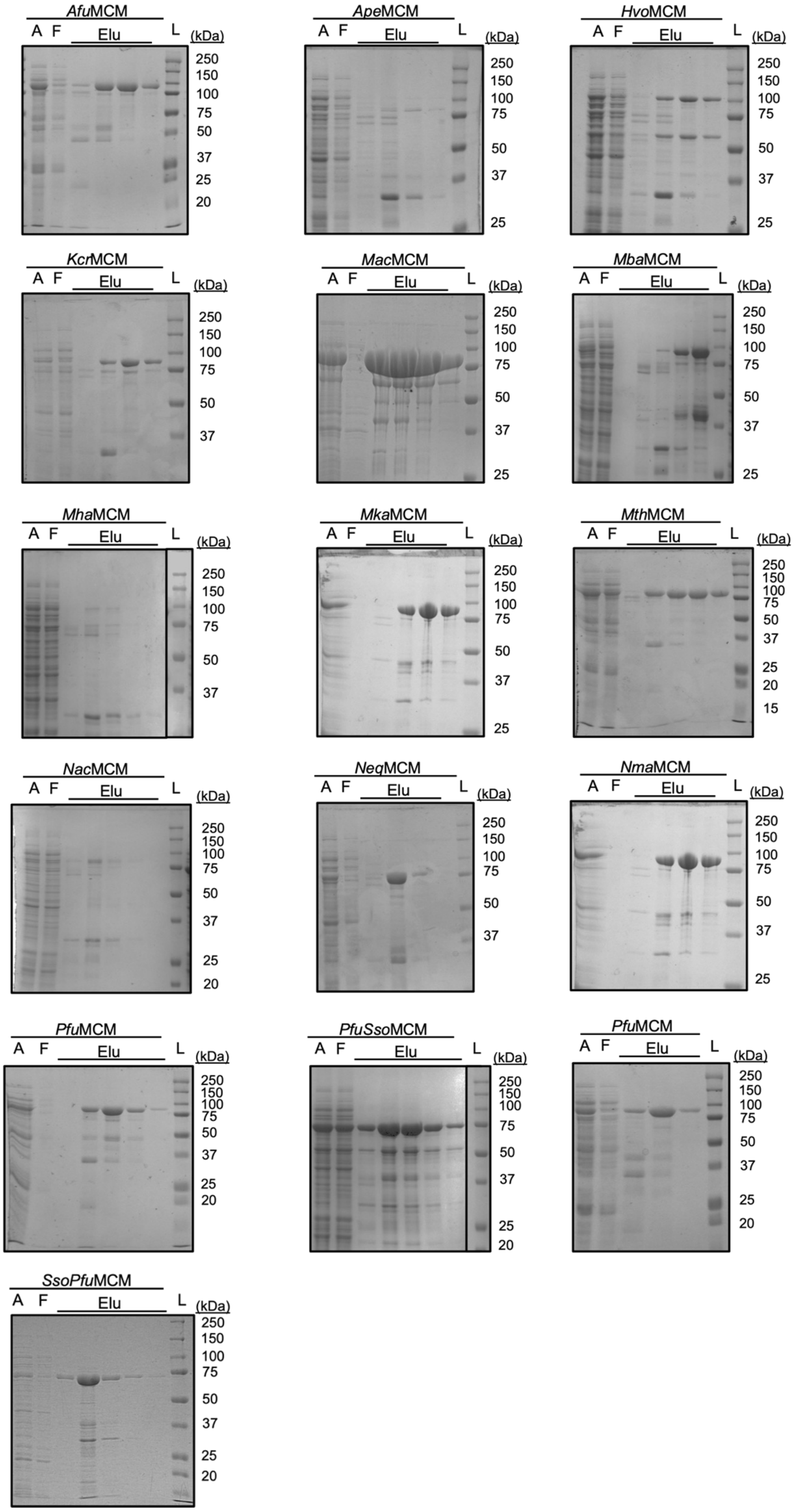


**Supplementary Figure 4: Purification of $His_{10}$-labelled recombinant MCM.** Fractions from IMAC purification were analyzed by SDS-PAGE on 12 % (*w/v*) polyacrylamide gels. Fractions include the sample applied to the Ni-NTA column (A), the column flow through (F) and select elutions (Elu). Reference (L) lanes represent MW standard marker (Precision Plus Protein™ All Blue Pre-Stained Protein Standards).

**Supplementary Table 2: ‘Crude’ purification yields and DNA contamination.**

| Identifier | Raw Yield (mg) | $A_{260/280}$ |
|---|---|---|
| *Afu*MCM | 4.14 | 0.83 |
| *Ape*MCM | 0.99 | 1.01 |
| *Hvo*MCM | 1.17 | 1.03 |
| *Kcr*MCM | 3.46 | 0.95 |
| *Mba*MCM | 1.20 | 0.92 |
| *Mha*MCM | 0.09 | 0.87 |
| *Mac*MCM | 18.4 | 0.63 |
| *Mka*MCM | 3.15 | 0.76 |
| *Mth*MCM | 5.65 | 0.82 |
| *Nac*MCM | 0.05 | 1.24 |
| *Neq*MCM | 1.54 | 0.76 |
| *Nma*MCM | 3.43 | 0.86 |
| *Pfu*MCM | 2.30 | 0.82 |
| $Pfu_N Sso_C$MCM | 2.39 | 0.71 |
| $Sso_N Pfu_C$MCM | 2.25 | 0.63 |
| *Sso*MCM | 2.80 | 0.88 |

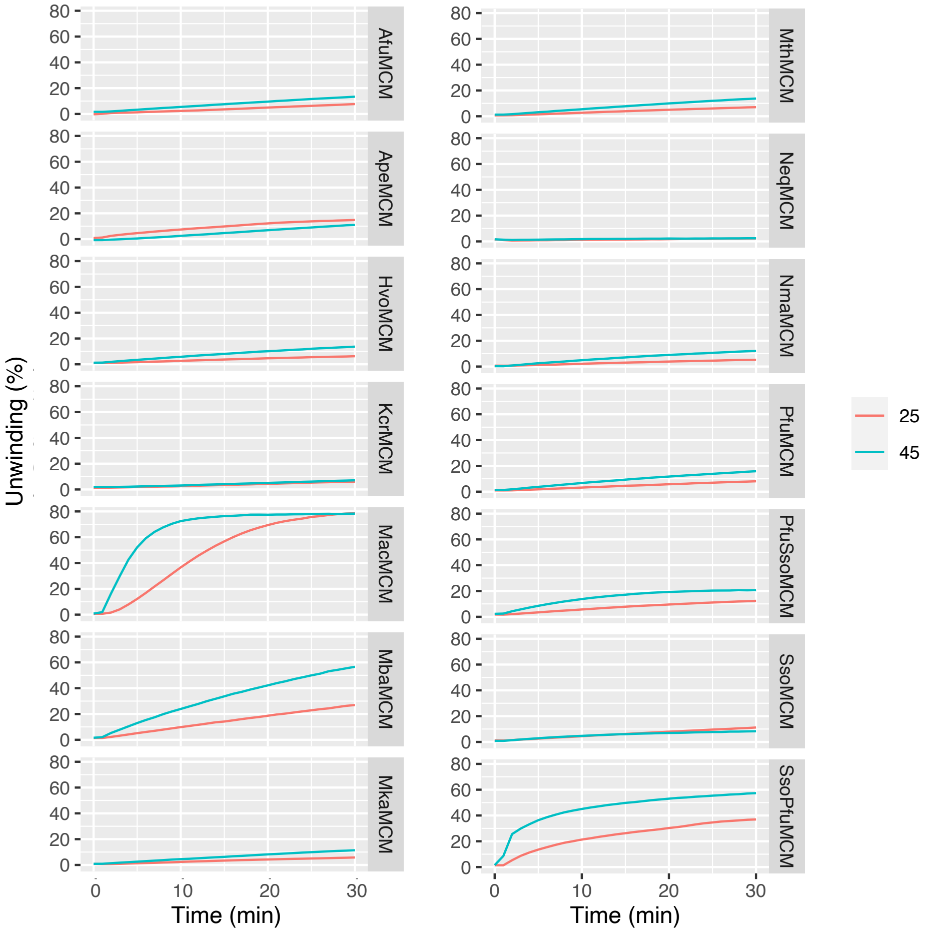


**Supplementary Figure 5: Real time unwinding traces for crude screens**.
Real time DNA-unwinding curves for the stated MCMs measured at 25 °C (orange) and 45 ºC (cyan). The slight sigmoidal shape seen for *SsoPfu*MCM was not evident when a pure sample was assays (see Supplementary Figure 11)

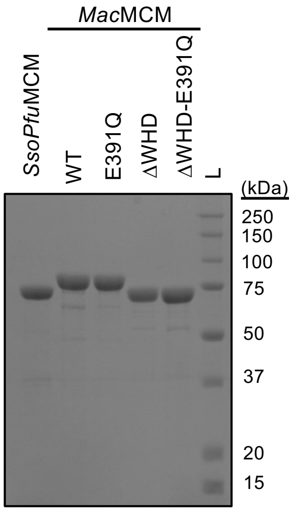


**Supplementary Figure 6: Purification of His$_{10}$-labelled recombinant MCM.** Purity analysis of purified recombinant proteins used in this study. SDS-PAGE analysis of 3 µg each purified MCM. WT, Wild-type; WHD, winged-helix domain.

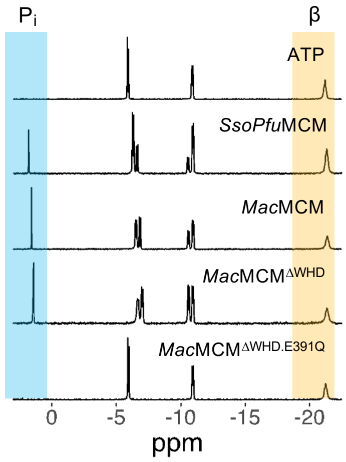


**Supplementary Figure 7: $^{31}$P-NMR confirms inactivity of ATPase mutants**
The presence of inorganic phosphate was determined for the stated MCM sample after 30 minutes using $^{31}$P-NMR. The top panel represents 50 mM ATP sample, without MCM. In each experiment, a final concentration of 8.3 µM MCM was added to ATP. The blue region highlights the expected region for the inorganic phosphate peak ($P_i$), whilst the orange region highlights the expected region for the β-phosphate of ATP.

| **Supplementary Table 3 Crystallographic statistics** | |
|---|---|
| **Data collection** | |
| Space group | C 1 2 1 |
| Cell dimensions | |
| *a,b,c* (Å) | 228.42 127.29 176.75 |
| *α, β, γ* (˚) | 90.00 91.71 90.00 |
| $R_{meas}$ | 0.115 (4.564) |
| I/σ <I> | 11.3 (0.3) |
| Completeness (%) | 99.99 (98.2) |
| Multiplicity | 6.8 (7.0) |
| $CC_{1/2}$ | 1.0 (0.5) |
| **Refinement** | |
| Resolution (Å) | 2.59 - 57.15 (2.59 – 2.64) |
| Unique reflections | 157088 (15547) |
| $R_{work}/R_{free}$ | 0.231 / 0.253 |
| No. of atoms | |
| Macromolecules | 26931 |
| Ligands | 176 |
| Solvent | 13 |
| *B*-factor (Å$^2$) | |
| Macromolecules | 96.43 |
| Ligands | 101.79 |
| Solvent | 67.64 |
| R.m.s deviations | |
| Bond lengths (Å) | 0.018 |
| Bond angles (º) | 1.49 |

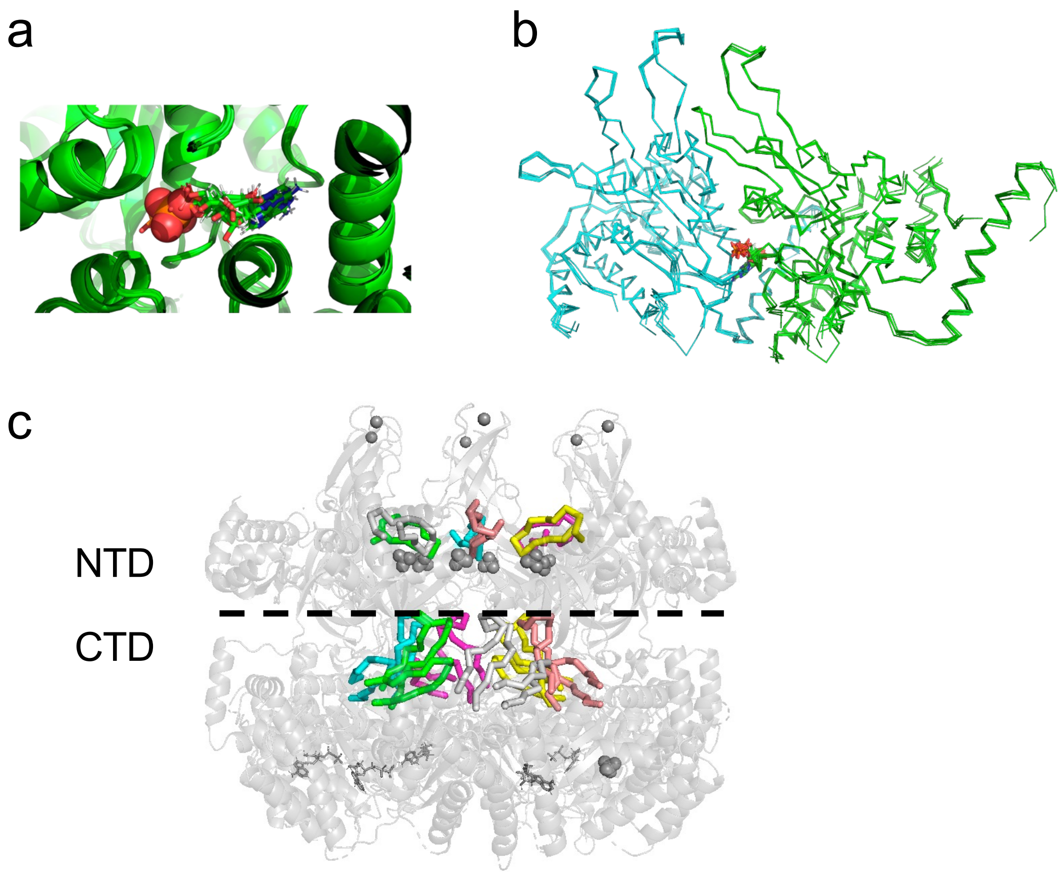


**Supplementary Figure 8: C-terminal domains are structurally indistinct.**
**a** View of structurally aligned ATPase active sites. Protein is represented in cartoon format. Ligands are represented in sphere (phosphate) and stick (ADP) formats. The phosphate is highlighted to emphasise that it mimics the position of the beta-phosphate of ADP. **b** View of structurally aligned ATPase active site pairs, visualized in ribbon format. This *cis*-acting subunit is coloured in cyan, whilst the *trans*-acting subunit is coloured in green. **c** Positioning of DNA-binding hairpins with respect to the N and C-terminal tiers of the *Mac*MCM hexamer (grey). Hairpins are displayed in ribbon format and coloured by subunit.

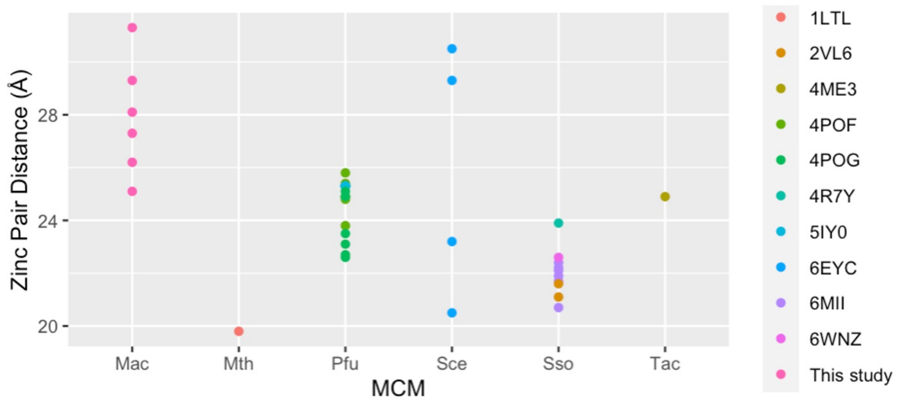


**Supplementary Figure 9: The distance between neighbouring Zinc fingers in *Mac*MCM.**
The distance between neighbouring *Mac*MCM Zinc Fingers was compared against structures published for archaeal and eukaryotic MCMs. Distances measured pertain to the position of the zinc-ion relative to the position of the zinc-ion on the neighbouring subunit. .

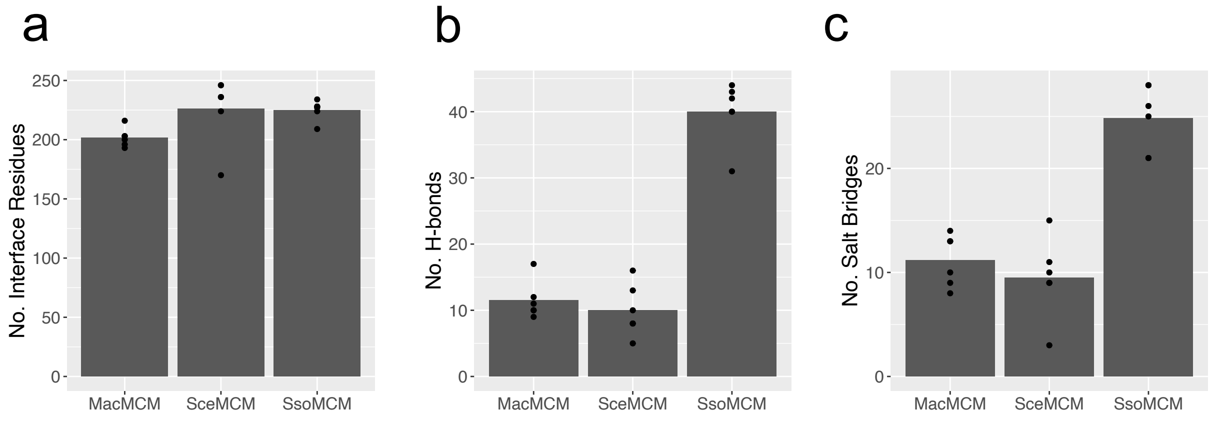


**Supplementary Figure 10: The interface composition of MCM hexamers from mesophilic organisms have similar properties.** Structures were imported into the PDBePISA (Proteins, Interfaces, Structures and Assemblies) portal and analysed. Data were then exported and averaged for each metric and MCM. *Mac*MCM (this study), *Sso*MCM (PDB: 6MII)[16], *Sce*MCM (PDB:6EYC)[17]. Black points represent raw subunit-subunit values. **a** Average number of residues involved in each interface. **b** Average number of hydrogen bonds formed at each interface. **c** Average number of salt bridges formed between MCM interfaces.

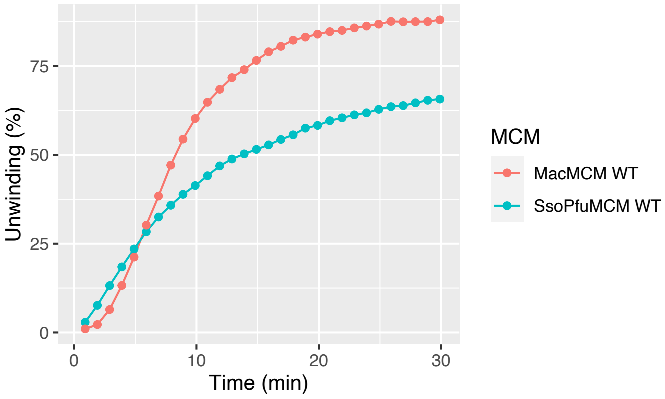


**Supplementary Figure 11: DNA unwinding kinetics of pure *SsoPfu*MCM.**
Real time DNA-unwinding curves for purified samples of *Mac*MCM and *SsoPfu*MCM at 25 °C.

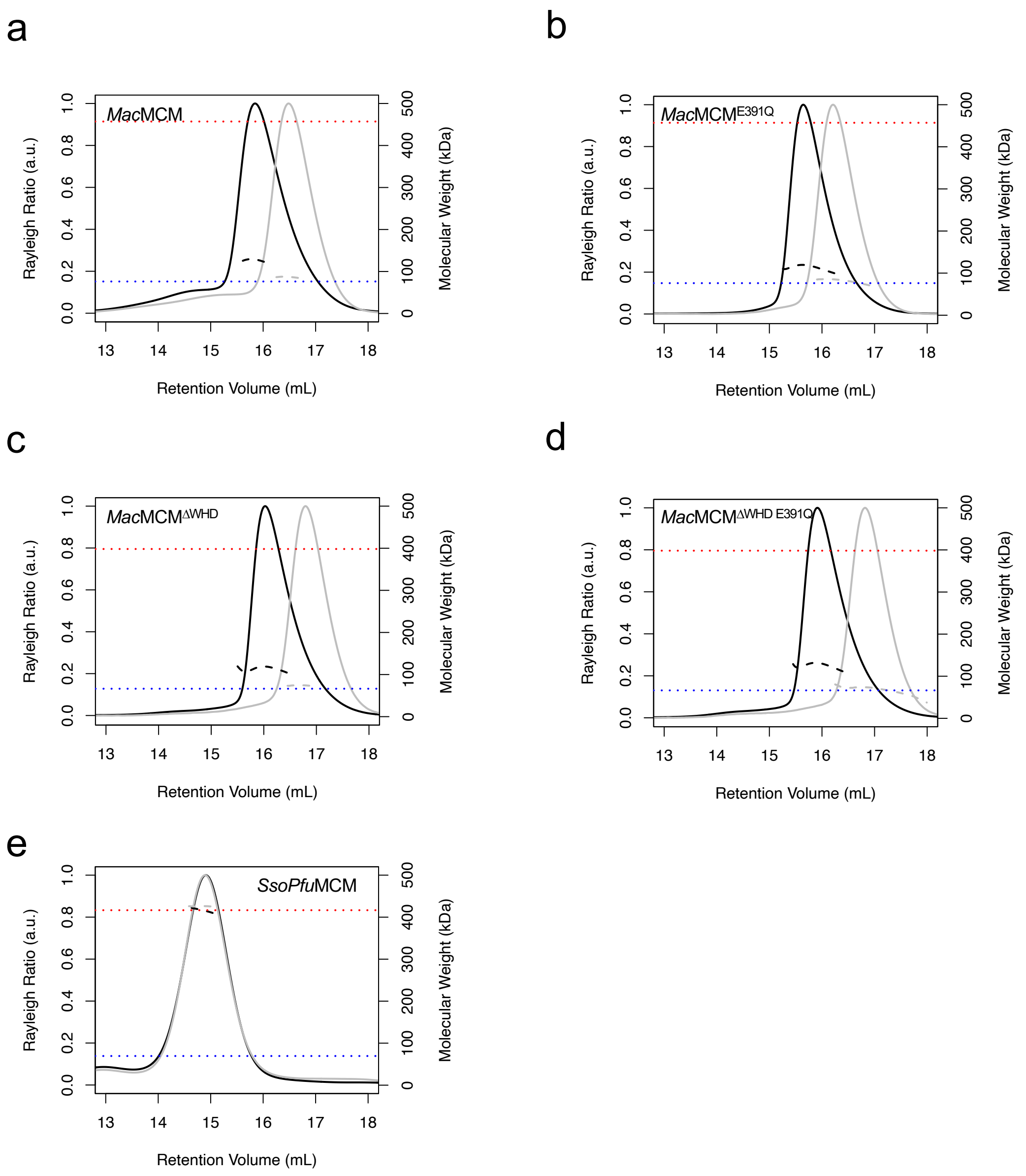


**Supplementary Figure 12. Oligomeric state of MCM samples in the absence of ligands.** **a-e** SEC MALLS elution profile of the stated MCM at concentrations of 1 mg/mL (grey; ~2 μM) or 10 mg/mL (black; ~20 μM) from a Superose 6 Increase column was monitored through light scattering (Rayleigh ratio). The Rayleigh ratio was normalised to the height of the main peak. Calculated molar mass values are shown as a dotted line. For each sample, expected molar masses are shown for a monomer (blue) and a hexamer (red).

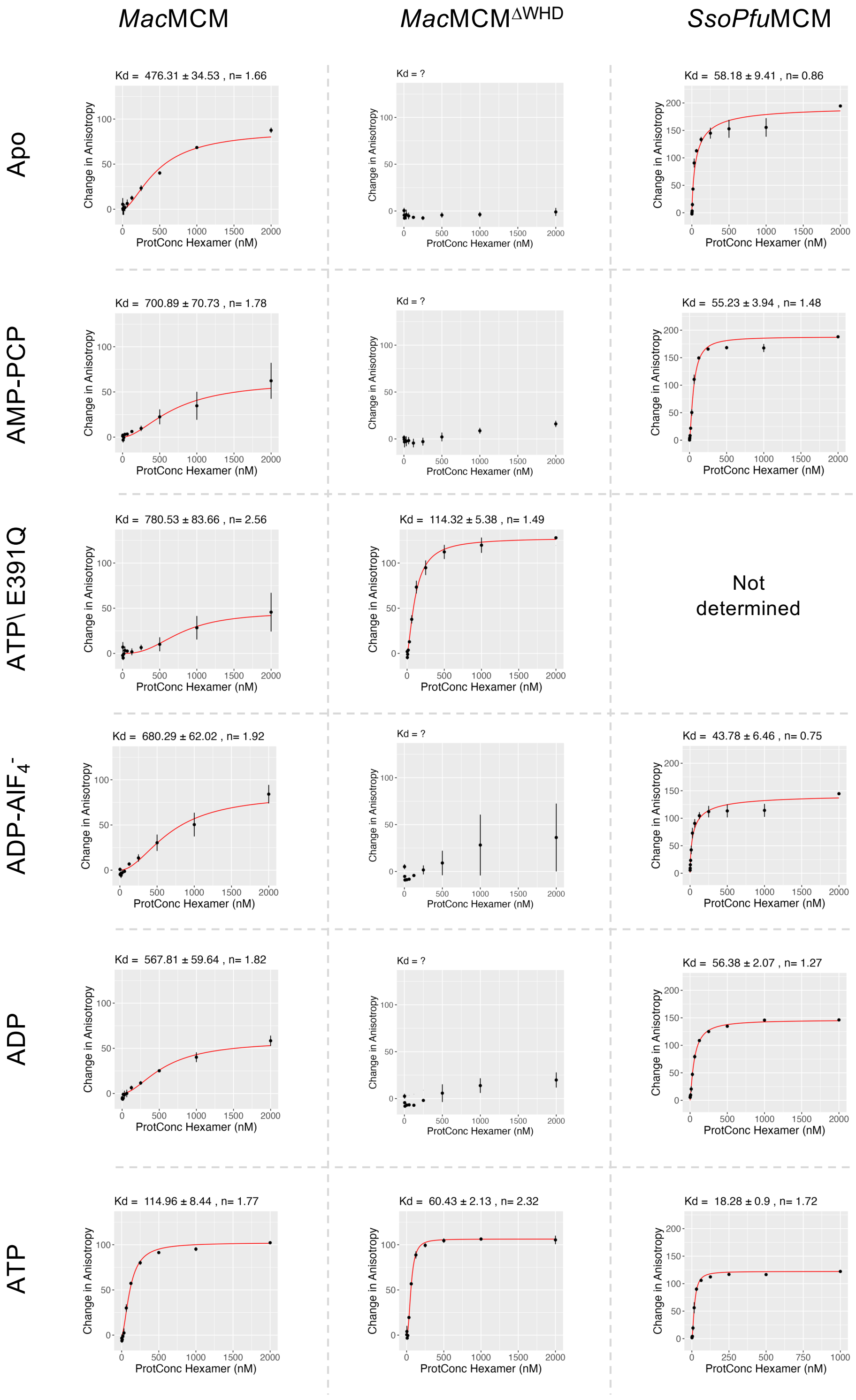


**Supplementary Figure 13. Assessment of MCM-DNA interactions in response to different nucleotides.** The binding of MCM to a forked DNA substrate was measured by fluorescence anisotropy. MCM were mixed at the stated concentration (nM hexamer) with 1 nM FAM-labelled forked DNA substrate and incubated for 30 minutes at 25 °C. Measurements were performed on a Clariostar plate reader (BMG Labtech). Anisotropy values were standardized by subtraction of a no-protein control well and fitted to Langmuir binding isotherm with Hill coefficient. Error bars represent +/-1 standard error of the mean, where n=3.

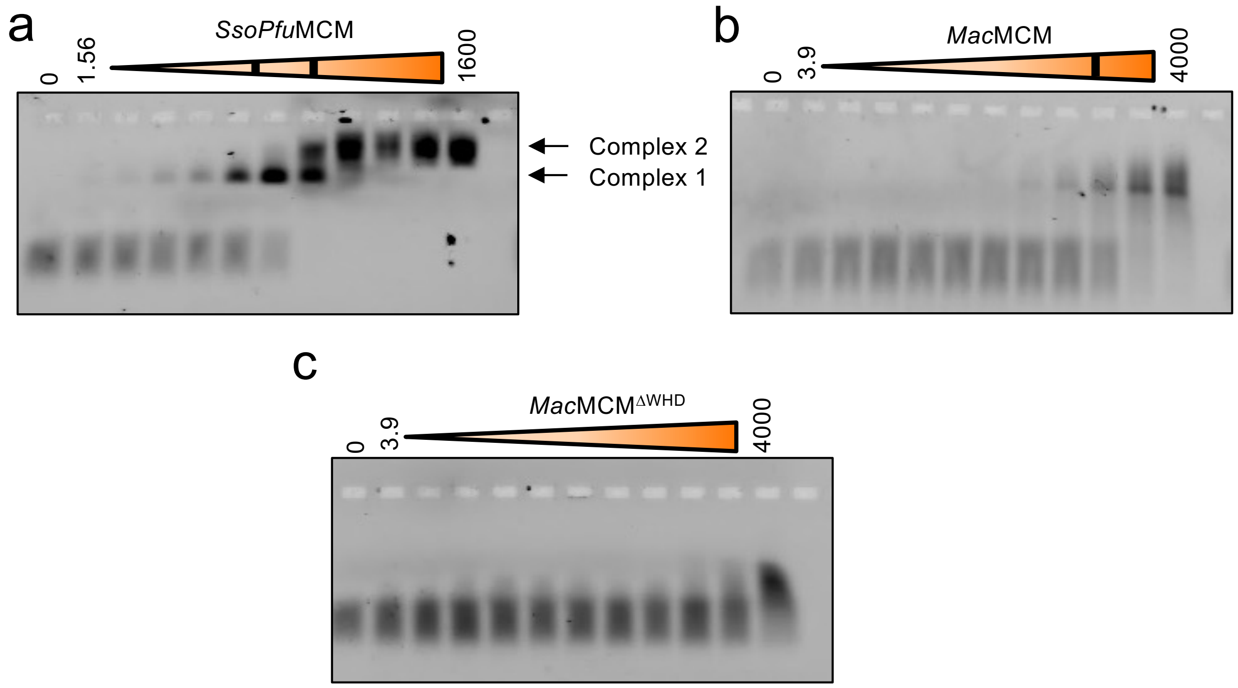


**Supplementary Figure 14. Assessment of DNA binding by EMSA.**
**a-c** MCM DNA binding was measured by EMSA. 10 nM fluorescein-labelled forked DNA substrate was mixed with different concentrations of each MCM and incubated for 30 minutes at 25 ºC Samples were resolved on a 1 x TB 0.8 % agarose gel and imaged using a Typhoon gel scanner. Horizontal triangles indicate maximum and minimum protein concentrations used in titrations with intermediate concentrations produced by 1:2 serial dilution.
